# Haplotype-based inference of recent effective population size in modern and ancient DNA samples

**DOI:** 10.1101/2022.08.03.501074

**Authors:** Romain Fournier, David Reich, Pier Francesco Palamara

## Abstract

Individuals sharing recent ancestors are likely to co-inherit large identical-by-descent (IBD) genomic regions. The distribution of these IBD segments in a population may be used to reconstruct past demographic events such as effective population size variation, but accurate IBD detection is difficult in ancient DNA (aDNA) data and in underrepresented populations with limited reference data. In this work, we introduce an accurate method for inferring effective population size variation during the past ~2,000 years in both modern and aDNA data, called HapNe. HapNe infers recent population size fluctuations using either IBD sharing (HapNe-IBD) or linkage disequilibrium (HapNe-LD), which does not require phasing and can be computed in low coverage data, including data sets with heterogeneous sampling times. HapNe showed improved accuracy in a range of simulated demographic scenarios compared to currently available methods for IBD-based and LD-based inference of recent effective population size, while requiring fewer computational resources. We applied HapNe to several modern populations from the 1, 000 Genomes Project, the UK Biobank, the Allen Ancient DNA Resource, and recently published samples from Iron Age Britain, detecting multiple instances of recent effective population size variation across these groups.

## 2 Introduction

The increasing availability of high-quality genomic data for both modern and ancient samples is creating exciting new opportunities for data-driven investigation of key evolutionary parameters. Among these, the effective size of a population plays an essential role in population biology^1^. A population’s effective size is defined as the number of individuals in an idealized evolutionary model^2,3^, and the ability to infer it from genomic data has a wide range of applications, including the study of past demographic events^4,5^ and cultural practices^6^, the quantification of the effectiveness of natural selection^1,7^, and the prediction of viability in conservation biology^8^.

Several statistical tools have been developed to reconstruct the trajectory of effective population size from genomic data^9^, each leveraging different genomic features and enabling the analysis of different data types. Methods that rely on the site frequency spectrum (SFS) of a sample^10–13^ avoid modeling recombination and are thus scalable, but require high-quality sequencing data to estimate the SFS and have been observed to be statistically inefficient^14^. Methods that model both mutation and recombination processes^15–19^, on the other hand, tend to scale to smaller sample sizes and require high-quality genome sequencing data. Recent approaches enable simultaneous modeling of recombination and allele frequencies in unphased sequencing data^18^, or scaling to larger sample sizes for accurately phased sequencing data^20^. Finally, several methods that focus on capturing the signature of recombination through the sharing of identical-by-descent (IBD) haplotypes^21–25^ or linkage disequilibrium^26–29^(LD) have been developed.

Inference of recent population size fluctuations is particularly appealing because it provides unique insights into demographic and evolutionary processes that are specific to the analyzed population. IBD-based methods have been used to infer recent demographic history^21–23, 25^ in SNP array and sequencing data. A key limitation of these methods is that they rely on accurate detection of IBD regions^30–33^. The performance of these algorithms depends on accurate long-range computational phasing, which may be hard to obtain, particularly in low coverage ancient DNA data. While being a less direct measure of the signature of past recombination events, LD-based summary statistics can be computed in unphased samples, including SNP array and ancient DNA data. LD has been extensively modeled^34–38^ and applied to infer effective population size^26–29, 38, 39^. The most recent methods for IBD- and LD-based inference, IBDNe^25^ and GONE,^29^ enable inference of population size fluctuations in time, without assuming a strictly parametrized demographic model. This strategy, however, poses additional challenges, due to the need to adequately regularize the inferred models^23,25^ to avoid reporting spurious fluctuations, while preserving manageable computational costs.

Here, we present a new method, called HapNe, that enables flexible inference of recent effective population size fluctuations using IBD or LD summary statistics, and can be used to analyze both phased and unphased SNP array or sequencing data, including low coverage or ancient DNA data with heterogeneous sampling time. Using extensive coalescent simulations, we show that HapNe accurately and efficiently infers recent demographic history, while regularizing the model to control for spurious oscillations in recent generations. We applied HapNe to reconstruct recent demographic history in both modern and ancient data, including populations from the 1,000 Genomes Project and different postcodes from the U.K. Biobank data set, where we observed a bottleneck in the Late Middle Ages corresponding to the period of the Black Death. We also analyzed ancient individuals from the Caribbean, Scandinavian Vikings, and individuals who lived in England during the Iron Age, observing isolation and expansion events that are consistent with past historical events, such as the transition from the Archaic to the Ceramic culture in the Caribbean.

## 3 Results

### 3.1 Overview of the HapNe algorithm

The HapNe algorithm infers recent effective population size using either IBD or LD data (see Methods and Supplementary Note for a detailed description of the algorithm). We refer to these two approaches as HapNe-IBD and HapNe-LD, respectively. HapNe-IBD uses IBD sharing information to compute summary statistics related to the count of IBD segments of different lengths. However, accurate detection IBD segments typically relies on phasing information and modeling of haplotype sharing to differentiate between identical-by-state (IBS) and truly IBD regions. Accurate phasing and haplotype modeling may not be possible if the analyzed genomes are not of high quality or not well represented in reference panels. HapNe-LD, on the other hand, leverages summary statistics related to long-range LD (Pearson correlation between sites). These LD statistics are easy to compute and do not require genotypes to be either phased or of high quality, enabling the analysis of past demographic events in low coverage or aDNA data.

HapNe-IBD and HapNe-LD both optimize a composite likelihood. To ensure that the model is appropriately regularized, HapNe utilizes a prior on the effective population size *N_e_*(*t*) that favors models with minimal population size fluctuations. When the analyzed IBD or LD data does not contain sufficient signal, this regularization mechanism prevents inferring spurious variation in *N_e_*(*t*), which may be incorrectly interpreted as past demographic events. The resulting approximate posterior is optimized to compute a maximum-a-posteriori (MAP) estimator of *N_e_*(*t*) and bootstrap resampling is used to provide estimates of uncertainty through approximate 95% confidence intervals. Both methods automatically exclude genomic regions harboring unusually large amounts of IBD or LD, which may be caused by natural selection or the presence of structural variation rather than past demographic events. In addition, HapNe-LD implements a test to detect the presence of possible biases due to the presence of strong LD caused by past admixture events (admixture LD) and can handle samples originating from different time points. The HapNe program is freely available as an open-source software package (see Code Availability).

### 3.2 Performance on simulated modern data

We used extensive coalescent simulations to benchmark HapNe-IBD and HapNe-LD against other recent methods for haplotype-based inference of recent effective population size. To this end, we considered several demographic scenarios (Figure 1a, dotted black lines), including: a constant population size of *N_e_*(*t*) = 20,000; an exponentially expanding population with 200,000 haploid individuals at *t* = 0 and 20,000 at *t* = 50 generations; an exponentially collapsing population with 2,000 living individuals at *t* = 0 and 20,000 at *t* = 100; and a population undergoing a strong bottleneck, evolving from 200,000 haploid individuals at *t* = 0 to 2,000 at *t* = 25, and then growing back to 20,000 at *t* = 50. For each of these populations, we simulated 256 diploid individuals. We generated realistic SNP-array data and used the simulated ancestral recombination graph to extract ground truth IBD segments longer than 1cM (see Methods).

**Figure 1:**
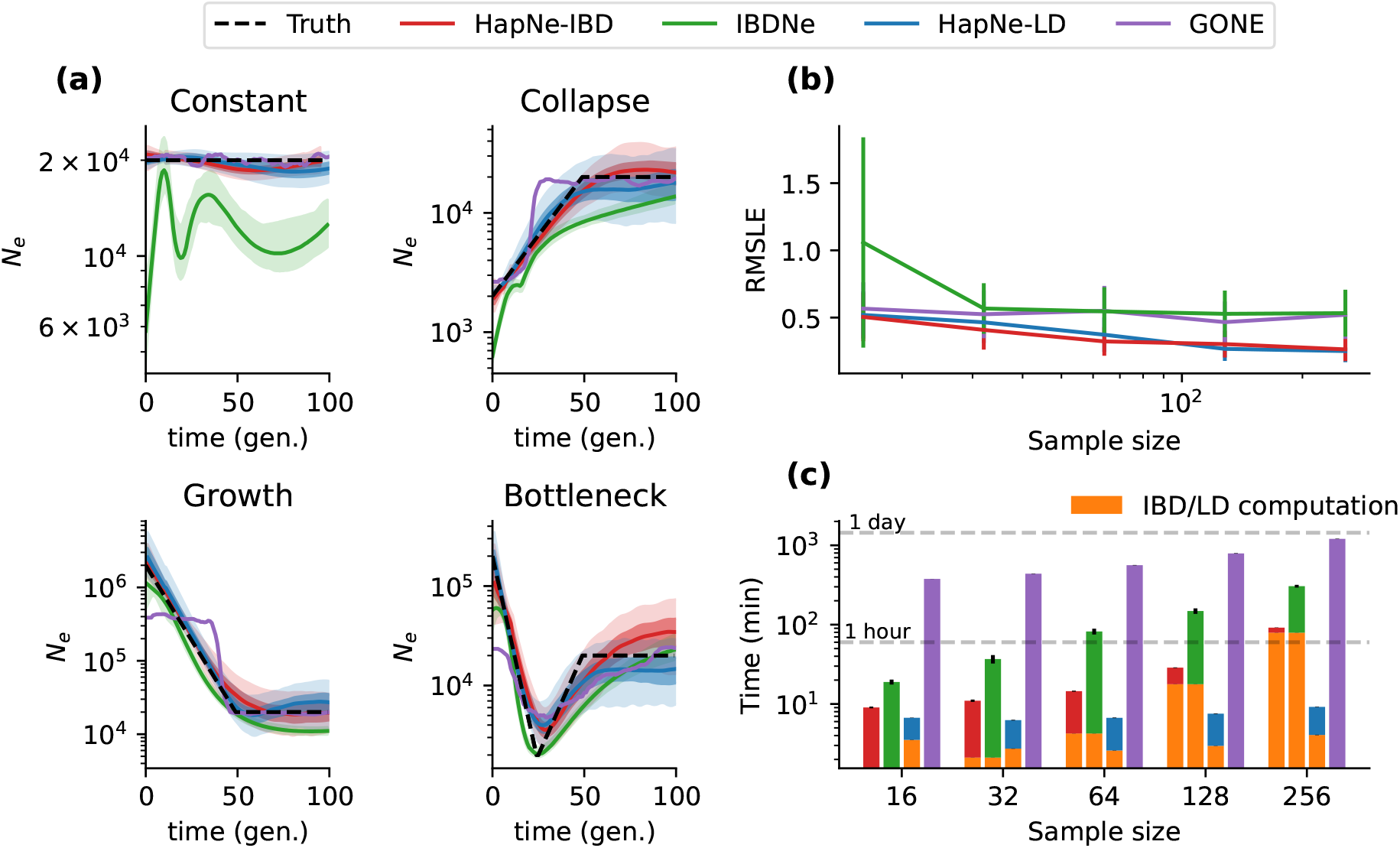
Benchmarks in simulated modern populations. (a) Comparison of HapNe-IBD, IBDNe, HapNe-LD, and GONE on simulated SNP-array data (256 individuals) for four different demographic scenarios. (b) Accuracy of the different methods on the ”Bottleneck” demographic model as a function of sample size. Error bars correspond to 1.96 × *SE* computed using 10 independent simulations. (c) Total running time for each method (including IBD segment detection and within-chromosome LD estimation, see Methods).

We initially considered the performance of HapNe-IBD and IBDNe^31^ in an idealized setting where ground truth IBD sharing information is available (see Supplementary Figure S1). In this scenario, HapNe-IBD generally produced lower error than IBDNe, measured using the root mean squared log-error (RMSLE) over the past 50 generations (see Methods). HapNe-IBD produced stable estimates of effective population size in the very recent past, whereas IBDNe tended to output spurious oscillations, a caveat that was highlighted by the authors^31^. We next inferred and analyzed LD summary statistics from the simulated array data using HapNe-LD. Because the LD signal reflects the presence of underlying IBD segments (see Supplementary Note), analysis of ground truth IBD data may be seen as an upper bound on the accuracy of HapNe-LD. We observed the RMSLE of HapNe-LD applied to SNP array data to be close to that of HapNe-IBD using ground truth IBD data, suggesting that HapNe-LD achieves close to optimal performance in these simulations, despite not utilizing phasing information (see Supplementary Figure S1b). We also tested the performance of GONE^29^, a recent LD-based method, and observed larger RMSLE in the past 50 generations (see Figure 1b). Due to its regularization procedure, HapNe-LD tended to infer smooth changes in population size, whereas GONE inferred more rapid fluctuations (see Figure 1a). GONE did not produce bootstrap confidence intervals in these simulations, due to an insufficient number of available SNPs (see Methods).

We next considered a more realistic scenario for the application of IBD-based methods (HapNe-IBD and IBDNe), where we inferred IBD sharing from simulated SNP array data (assuming perfect phasing, see Methods). We detected IBD sharing using the FastSMC program^32^; similar results for IBDNe were obtained by using the recommended HapIBD software^33^ (see Supplementary Figure S2). Figure 1a shows the output of all four methods on a data set of 256 diploid samples and results for other sample sizes are summarized in Figure 1b (also see supplementary figures S3 and S4). In most cases, the noise introduced by inferring IBD from the data resulted in biases in the inferred effective population sizes; IBDNe tended to underestimate recent effective population size, while HapNe-IBD tended to overestimate ancestral population size (Supplementary Figure S3). We observed the error in IBD detection to be dependent on several factors, including demographic history and the length of the inferred segments (see Supplementary Figure S5). We note that additional biases due to genotyping and phasing errors are likely to be present in real data, further affecting the quality of IBD-based analyses.

We finally benchmarked the computational speed of these methods and observed HapNe-IBD and HapNe-LD to be more computationally efficient than IBDNe and GONE (see Figure 1c). Computing LD scales only linearly with the number of analyzed samples, while detecting pairwise IBD sharing requires computation that is quadratic in the number of samples, making LD-based analyses more scalable. Unlike IBDNe, which requires more time to fit larger samples, HapNe-IBD only computes a fixed-size vector of the IBD segment lengths, significantly reducing computational costs for larger samples. The difference in computational time between HapNe-IBD and HapNe-LD is mainly driven by differences in the time required to compute IBD and LD summary statistics.

Overall, HapNe-IBD and HapNe-LD provided improved accuracy and substantially reduced computational times compared to existing methodologies. Although IBD-based inference of effective population sizes is potentially more accurate than LD-based analysis, the need to accurately detect IBD sharing is likely to introduce substantial biases in the inferred population sizes. HapNe-LD’s performance was observed to be close to that of IBD-based methods applied to ground truth IBD data and may be applied in the analysis of large sample sizes, providing several practical advantages over IBD-based methods in the analysis of real data sets.

### 3.3 Performance on simulated aDNA data

HapNe-LD does not require phased or high coverage data, making it especially suitable for the analysis of effective population sizes of ancient populations, where phase determination can be poor. However, LD-based analysis suffers from several limitations and potential confounders, some specific to aDNA data. First, analyses based on aDNA data sets tend to contain fewer samples sequenced at relatively low coverage compared with modern panels. Furthermore, different sequencing strategies balancing sample size and coverage might lead to different performances in effective population size inferences. Next, an important confounder is the potential presence of admixture in the analyzed samples, which is often encountered in real populations as a result of past demographic interactions and induces long-range correlations among genomic variants^40^. Finally, individuals sampled at a site are unlikely to have lived at the same time, with a few notable exceptions^41,42^. If not modeled, this source of time heterogeneity may lead to biased effective size estimates.

We set out to test HapNe-LD’s robustness to these sources of confounding. We first created synthetic aDNA samples by generating pseudo-diploid individuals with different levels of missingness *m*, mimicking the effects of reduced sequencing coverage *C*, with *m* ≈ *e*^−*C*^ (see Methods). We tested the relative impact of the simulated sample size *s* and coverage on HapNe-LD’s inference accuracy (see Figure 2a and Supplementary Figure S6 for additional demographic scenarios). As expected, RMSLE decreases when more samples are available and when coverage increases (see Figure 2b and Supplementary Figure S7). We then tested whether HapNe-LD would perform better when analyzing a larger number of low-coverage samples rather than a smaller number of high-coverage samples. To this end, we performed simulations where the overall number of sequencing reads is kept approximately constant, while the number of analyzed samples and their coverage are varied (see Figure 2c and Supplementary Figure S7). We considered an analysis involving 256 individuals and observed that reducing coverage from 30x to 1.4x had no significant impact on the performance while requiring only about 5% of the reads. Using an equivalent number of reads to perform high coverage (30x) sequencing would only allow sequencing 16 individuals, resulting in significantly higher RMSLE. These results suggest that sequencing at a coverage higher than 1-2x does not lead to significant improvements in HapNe-LD’s performance, and that HapNe-LD is more accurate when a larger number of individuals is sequenced at lower coverage compared to settings in which a smaller number of high coverage samples is analyzed.

**Figure 2:**
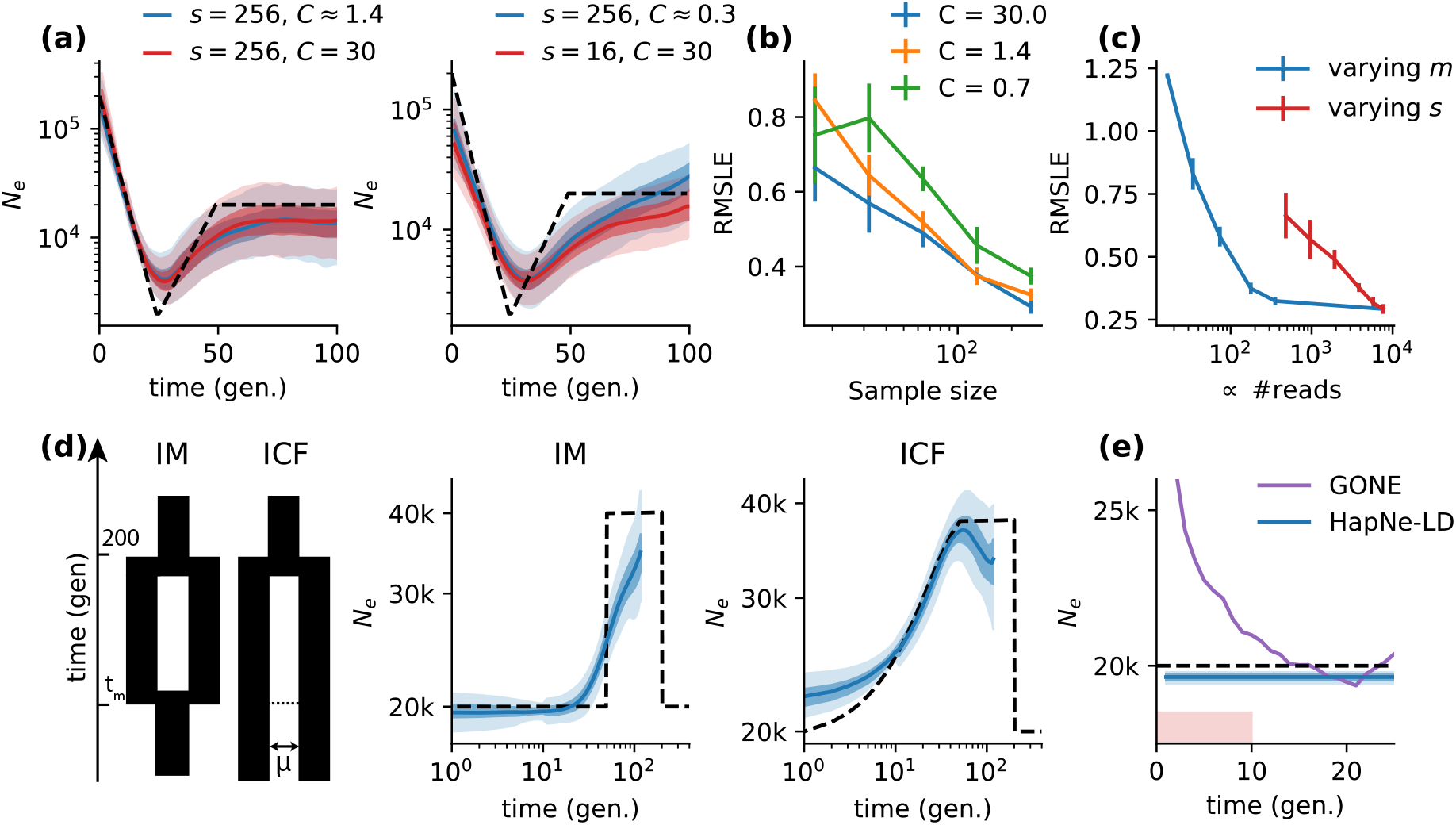
Results in simulated aDNA data. **(a)** HapNe-LD inference results for simulated aDNA-like data under the ”Bottleneck” demographic scenario (dashed lines) where the number *s* of simulated samples and fraction *m* of missing SNPs, or equivalently the coverage *C*, are varied (see Methods). **(b)** RMSLE over the first 50 generations for different coverage levels. Error bars correspond to 1.96 × *SE* computed using 10 independent simulations. **(c)** Comparison of the accuracy of HapNe-LD based on two sequencing strategies. The red line reports RMSLE for high coverage data (*m* = 0, *C* = 30) with varying sample size *s*. The blue line reports RMSLE for fixed *s* = 256 and varying coverage. Error bars correspond to 1.96 × SE computed using 10 independent simulations. **(d)** HapNe-LD results under the IM and ICF models of recent admixture, depicted on the left. For both models, we set *t_m_* = 50 generations. For ICF simulations, we sampled all individuals from one population and selected a migration rate *μ* such that ancestors of a sampled individual are located in the second population with probability close to 1/3 (see Methods). **(e)** HapNe-LD and GONE inference results for a simulation where individuals from a population of constant size of *Ne* = 20,000 are uniformly sampled over an interval Δ*T* = 10 generations (red shaded area).

We next simulated a population affected by recent admixture (see Supplementary Note) by considering two demographic scenarios (similar to those used in ^43^). In these scenarios, two isolated populations first separate and then either merge again (IM model) or experience continuous gene flow (ICF model, see Figure 2c). All simulated models had a constant number of 20,000 haploid individuals within each population; the interaction time *t_m_* was set to 50 generations. Simulation results for other values or *t_m_* are shown in Supplementary Figure S8. For the ICF model, we sampled all individuals from one population and selected a migration rate *μ* such that at time *t_m_* the ancestral lineages of all individuals are located in the second population with a probability close to 1/3. Figure 2c shows that HapNe-LD results under these models do not strongly deviate from the true underlying effective population size (see Supplementary Note). Some ICF simulations resulted in an increase in the inferred recent population size (see Supplementary Figure S8), likely due to model regularization, indicating that larger sample sizes are needed to infer subtle population size variation at these time scales. Taken together, these results suggest that HapNe-LD is robust to reasonable levels of admixture LD. The HapNe-LD software implements a statistical test for admixture LD, warning the user if significant admixture LD is detected.

Lastly, we considered potential biases arising due to heterogeneous sampling times of the analyzed aDNA individuals. We used analytical modeling (see Methods and Supplementary Note) to confirm that, if not accounted for, heterogeneous sampling times lead to biased recent effective population size estimates. We performed simulations of aDNA samples originating from heterogeneous time locations under a constant demographic history, uniformly drawing the time offset of each sample between 0 and Δ*T* generations in the past (see Methods). In this setting, we observed that using GONE to infer effective population size leads to the spurious inference of a recent population expansion, consistent with analytical predictions under unmodeled time heterogeneity (see Figure 2d). The HapNe-LD algorithm allows utilizing prior knowledge of sampling times (e.g. from radiocarbon dating or archeological context) in the form of a user-provided time interval for each analyzed individual (see Methods). Using simulations, we verified that this approach effectively removes recent biases due to time heterogeneity.

### 3.4 Inference of recent effective population sizes in the UK Biobank and 1,000 Genomes Project data sets

We used HapNe-IBD and HapNe-LD to analyze recent effective population size variation within the UK Biobank data set. Accurate inference of recent demographic events requires a combination of large sample sizes and small effective population sizes, which make it possible to estimate recent coalescent rates. In this case, large recent effective population sizes generally present across the UK are balanced by the large sample sizes available in the UK Biobank data set. In order to mitigate the impact of admixture LD, we focused on the larger group of samples with self-reported white British ancestry, and only considered unrelated individuals to avoid biasing demographic inference in recent generations. We grouped individuals based on the postcode of their self-reported birthplace and report analyses for three of these postcodes (see Figure 3a, Methods). We also used FastSMC to detect IBD segments within each of these postcodes. Regions with unusually high LD or IBD sharing were excluded using HapNe’s filter (Supplementary Figure S9).

**Figure 3:**
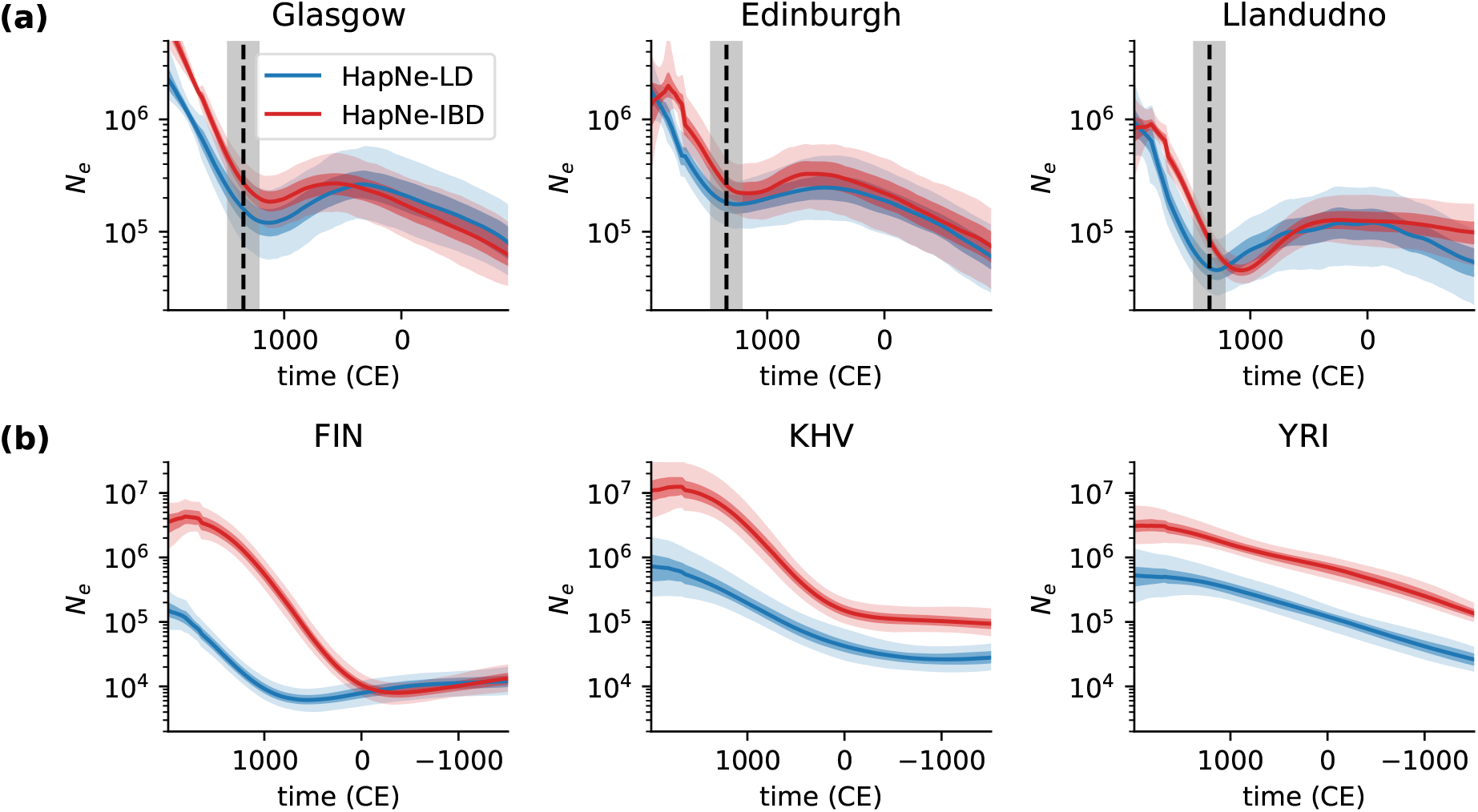
HapNe-IBD and HapNe-LD estimates of recent effective population sizes in modern populations. **(a)** Inference results for three postcodes: Glasgow (G), *s* = 14,724; Edinburgh (EH), *s* = 9,981; and Llandudno (LL), *s* = 2,089 from the UK Biobank data set. The vertical dashed line corresponds to the estimated date of the Black Death in the UK (1348,^44^). HapNe results are converted to years assuming 29 years per generation. The shaded grey area depicts how the placement of the Black Death would shift with respect to the inferred demographic models if values between 23 and 35 years per generations were assumed. **(b)** Inference results for three populations (Finnish, FIN, *s* = 99; Kinh in Ho Chi Ninh City, Vietnam, KHV, *s* = 99; Yoruba in Ibadan, Nigeria, YRI, *s* = 107) from the 1,000 Genomes Project.

Effective size trajectories inferred from these regions in the UK all exhibit a bottleneck event during the Late Middle Ages, which roughly corresponds to the period of the Black Death (Figure 3a, vertical dashed line). The inferred population size for individuals from the Llandudno postcode has a significantly smaller effective population size compared to the ones inferred for Glasgow and Edinburgh. Such a smaller effective size offers a stronger source of recent demographic signal, allowing to perform inference using a smaller sample size (*s* = 2,089 for Llandudno, *s* = 14,724 for Glasgow, and *s* = 9,981 for Edinburgh). In contrast, detecting the more subtle contraction to a larger minimum bottleneck size in Glasgow required a substantially larger sample size, as highlighted when we downsampled data from this postcode to 2,000 individuals (see Supplementary Figure S10). In this experiment, the bottleneck was only apparent in the output of HapNe-IBD, suggesting that LD-based analysis may lead to comparably lower statistical efficiency in cases where high-quality IBD signal is available. Demographic models inferred by HapNe-IBD and HapNe-LD are broadly consistent, although HapNe-IBD tends to report a larger effective population size, with a significative shift towards more remote times. These observations are compatible with the presence of underlying IBD segments that are undetected or broken into smaller segments, due to the presence of phasing or genotyping errors in the data.

We next applied HapNe-IBD and HapNe-LD to data from the 1,000 Genomes Project (1kGP,^45^). Unlike the UK Biobank, most 1kGP groups contain a small number of samples, which originate from large populations. Furthermore, several groups represented in the 1kGP data set are known to have undergone recent admixture, which complicates LD-based analyses^45^. We therefore expected analysis of recent effective population sizes to only be possible in a small subset of 1kGP populations. We used HapNe-LD to compute LD for each population and estimated recent IBD sharing using the FastSMC algorithm^32^ (see Methods). We used HapNe’s filters to exclude populations that were flagged as either not containing sufficient recent demographic signals or exhibiting strong admixture LD (19/26). We then inferred recent effective population sizes using the HapNe-LD and HapNe-IBD methods.

Figure 3b shows results for three populations that passed these filters. Results for all populations without significant admixture LD are shown in Supplementary Figure S11, which also reports results obtained by running the IBDNe algorithm. Supplementary Figure S12 shows two additional populations passing these filters for a less stringent significance cutoff and Supplementary Figure S13 displays the remaining 19 groups. Again, the demographic history inferred using IBD data consistently resulted in larger effective population sizes compared to LD-based results, particularly for recent generations, and were more strongly regularized due to reduced signal. These effects were more pronounced in these groups compared to the UK Biobank analysis, likely due to smaller sample sizes leading to lower phasing and IBD detection quality. HapNe-LD suggests a recent expansion for the individuals from the Kinh population in Ho Chi Minh City, Vietnam (KHV) and the Yoruba population in Ibadan, Nigeria (YRI) and infers a bottleneck at 1,000 CE for the FIN population, consistent with previous reports^25,29,46^. These demographic events are inferred to have an earlier onset using IBD data, likely also a result of noisy IBD detection. We also observed that IBD-based methods inferred strong bottlenecks in many African and South American populations around 1,000 CE, which is likely due to biases in the IBD-detection (see Supplementary Figure S13).

Overall, these results suggest that HapNe-LD and HapNe-IBD provide similar results when large samples and high-quality IBD data are available. HapNe-LD, however, provides more robust results than HapNe-IBD in data sets where phasing and IBD detection accuracy are reduced, at the cost of an only slightly reduced statistical efficiency. HapNe-LD may produce biased estimates for data sets including a history of strong recent admixture, as highlighted for some populations in Supplementary Figure S13. These biases usually result in an apparent population collapse in the recent past; in these analyses, however, HapNe-LD implements tests to flag populations where strong admixture is likely to result in such a spurious recent bottleneck.

### 3.5 Inference of recent demographic history in ancient populations

We applied the HapNe-LD method to aDNA sampled from four different sites for which large cohorts from similar time strata were available (see Methods and supplementary tables S1–S7).

We first analyzed a group of recently published individuals excavated in Pocklington, Yorkshire, UK^47^ (see Figure 4a). The archeological context suggests that this group belongs to the Arras culture, which is distinctive relative to other Iron Age cultures in the UK but shows similarities with contemporary cultures in the Paris Basin and Ardennes/Champagne regions of France. These individuals were found to be unusually highly drifted from nearby groups, although their F-statistics do not highlight significantly divergent admixture histories^47^. This suggests that these groups share common origins but may have been isolated for some time. To test this, we compared the effective population size for 24 individuals from the Arras culture to that of 49 other Iron Age individuals from Southern England (supplementary tables S2 and S3). For the Arras, we detected a significant recent population contraction, starting between 500 and 1,000 BCE, which was not observed in individuals from Southern England. This is consistent with isolation of the Arras group from other Iron Age individuals in the South of England, possibly also reflecting isolation by distance due to the stronger geographic localization for the Arras samples. Admixture LD for these groups was found to be negligible, suggesting that the observed demographic signature is not due to admixture (see Supplementary Table S1). The small population size of the Arras group might also explain why this population was found to be unusually highly drifted from nearby groups. The recent effective population size inferred for individuals in the South of England was compatible with population size estimates obtained for modern UK Biobank individuals, although confidence intervals were large over the first 1,000 years due to a reduced sample size.

**Figure 4:**
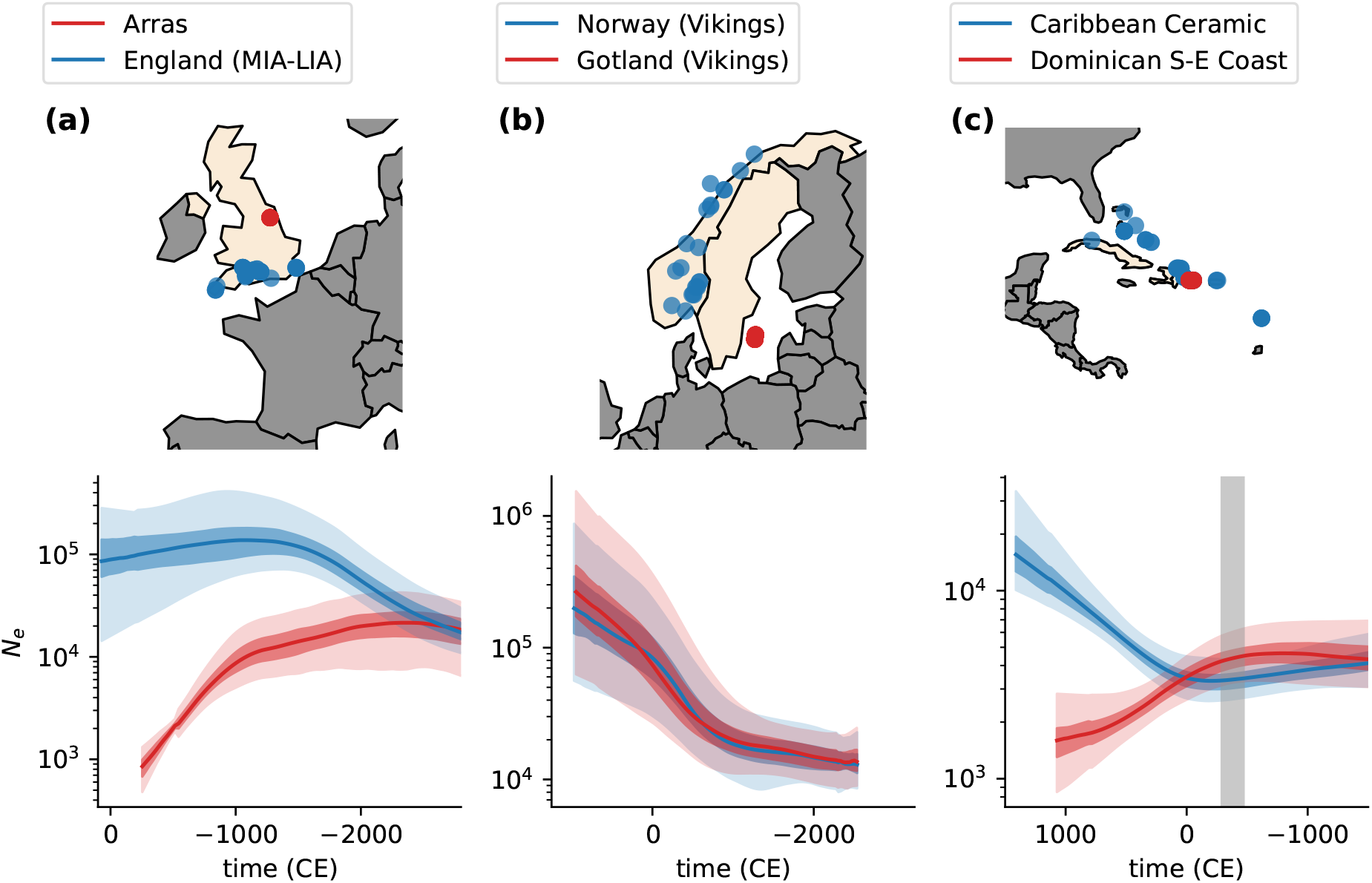
**(a)** Analysis of 49 Middle to Late Iron Age individuals from South England, compared to 24 individuals related to the Arras culture near Yorkshire. **(b)** Inference based on 22 Viking samples found in modern Norway (blue) and 28 found in Gotland, a Swedish island (red). **(c)** Effective population size inference based on 71 unrelated individuals from the Caribbean Ceramic clade and 18 from the Dominican South-East coast subclade. The grey shaded area corresponds to the estimated date for the transition from Archaic to the Ceramic culture in the region.

We next analyzed 22 genetically similar individuals from the Viking Age buried in Norway, together with 28 individuals from the south-east Swedish island of Gotland^41^ (Figure 4b and supplementary tables S4 and S5). Norwegian and Swedish Vikings have been observed to have a slightly smaller proportion of ancestry from Neolithic farmers from Anatolia compared to Swedish Vikings. On the other hand, Vikings from Gotland have a relatively higher estimated fraction of ancestry shared with Bronze Age individuals from the Baltic region. Despite these differences, the demographic histories inferred by HapNe-LD for the recent past of these individuals substantially overlap, and both trajectories show a significant expansion during the iron age (−500 to 800 CE).

Finally, we focused on 71 unrelated individuals from the Caribbean, first analyzed in Fernandes et al.^48^ (n=62) and Nägele et al.^49^(n=9) spanning ~1,149 to ~1,440 CE (supplementary tables S6 and S7). For these samples, HapNe-LD infers a weak sign of a bottleneck occurring around 1 CE, followed by a significant expansion, as shown in Figure 4c (blue line). This pattern may reflect the transition from the Archaic to Ceramic context about 2,500-2,300 years ago (Figure 4a, grey area), which has been associated with migration events in the region^48^. We also extracted and separately analyzed a subgroup of individuals from South-East Dominican sites (Figure 4c, red). These individuals are part of a subclade previously identified in^48^. The population size inferred for this group matches that of the broader Caribbean group in the deep past, consistent with common origins, but shows a distinctive sign of contraction in the more recent past. Admixture LD is detectable in these individuals, which may partially explain the observed contraction, as observed in some 1kGP populations (see Supplementary Figure S13 and Supplementary Table S1). Nevertheless, the sizes inferred by HapNe-LD in the recent past roughly match those inferred using runs of homozygosity^50^, supporting the possibility of a population contraction starting after the transition from the Archaic to the Ceramic Culture^48^. As in the case of the Arras and Southern England individuals, these demographic patterns may also be due to isolation by distance, where samples originating from different islands result in a larger effective size when considered together.

## 4 Discussion

We developed an algorithm, called HapNe, that leverages the count of IBD segments of different lengths (HapNe-IBD) or long-range LD (HapNe-LD) to infer recent effective population size fluctuations in modern or ancient DNA data. HapNe-IBD and HapNe-LD implement a number of preprocessing steps, as well as tests to verify that sufficient recent demographic signal is present in the data and to detect the presence of admixture LD. Both methods minimize a power-likelihood based on an analytic link between observed summary statistics and the effective population size and use regularization to avoid producing spurious oscillations. We used extensive simulation to show that both HapNe methods were more accurate and computationally faster than available algorithms for IBD-based and LD-based inference of recent demographic history, producing lower error and fewer spurious oscillations. These simulations also showed that while HapNe-LD does not require high quality or phased data and scales better with sample size, its performance can be close to that of IBD-based methods applied to ground truth IBD information. Finally, we applied HapNe to several modern and aDNA data sets, detecting evidence for recent past demographic events across these populations. These include population size contractions corresponding to the period of the Black Death in different regions of the UK, as well as bottleneck and expansion events in 1,000 Genome Project populations. In aDNA data, these analyses provided evidence for divergence and isolation events, as well as shared demographic histories in subgroups from several ancient populations with diverse geographic and temporal origins.

Our analyses suggest that LD-based inference of recent demographic variation provides a route to circumenting biases that may arise in IBD-based demographic inference. Although the spectrum of shared IBD haplotypes is an effective source of information for analyses of past demographic events, accurately estimating IBD sharing is complicated in low coverage and aDNA data and may lead to biased results. This may also be the case in modern populations when limited data availability prevents accurate phase estimation. Although summary statistics of LD rely on less direct observation of historical recombination events, they may be effectively computed in unphased and low coverage data sets. This enables analyzing recent demographic events in samples from poorly represented populations and, coupled with modeling of heterogeneous sampling time, in aDNA data sets. Performing both IBD-based and LD-based analyses may offer validation for an inferred demographic model and allow testing for the presence of biases in either approach. An additional source of potential bias in methods for demographic inference is linked to the need to make assumptions about the type of demographic model being inferred. In this context, approaches that avoid relying on a predefined set of models provide more flexibility, but require further tuning strategies to balance the desired sensitivity to past demographic events with the need to prevent the inference of spurious fluctuations. Our work suggests that the use of self-tuning regularization mechanisms helps mitigate the risk of spurious inferred fluctuations. Finally, our analyses highlight the importance of accurately preprocessing both IBD and LD signals before performing demographic inference, as results may vary significantly if unfiltered data is utilized. Key preprocessing steps include testing for the presence of admixture LD and systematically filtering out regions of the genome that harbor unusually high IBD sharing or LD (see e.g. Supplementary Figure S9). These may be due to natural selection or the presence of structural variation and lead to biases in analyses of demographic history and selection if not accounted for.

We outline several limitations and directions of future development for this work. First, HapNe-LD assumes that the LD signal observed in the data is solely due to past population size fluctuations. In some instances, residual admixture LD can be present in the data after filtering, causing a spurious bottleneck in the recent past and creating the need to carefully interpret models that resemble this type of signature. Similarly, HapNe-IBD currently only relies on the observed spectrum of IBD sharing, which may be biased due to inaccurate IBD detection. Future work may allow explicit modeling of type-1 and type-2 errors in IBD detection, mitigating biases in the inferred demographic models. Second, while regularization helps prevent the inference of spurious demographic fluctuation, it leads to favoring constant and exponential demographic histories that lack fluctuations if these are not supported by the data. When interpreting demographic models inferred by HapNe, it is important to note that an inferred constant growth rate may reflect insufficient evidence for past demographic variation (see e.g. Figure S10). Finally, HapNe-LD makes several model simplifications, including the assumption that the analyzed samples come from a single population. HapNe may be extended to explicitly account for multiple populations, improving the analysis of more complex demographic models such as those involving isolation by distance, divergence, and admixture. Similarly, HapNe-LD is currently focused on the inference of recent demographic history, but may be extended to the analysis of deeper time scales by modeling variation in allele frequencies, which are currently assumed to be constant in time. Despite these limitations, we expect that the HapNe framework developed in this work will offer valuable insights into past demographic events in both modern and ancient DNA data.

## 5 Methods

### 5.1 Simulated genetic data

We used the ARGON simulator^51^ (version 0.1.160415) to generate synthetic genotypes and ground truth IBD data for modern and ancient populations. Simulations with time heterogeneity were performed using msprime^52^ (version 1.1.1). We simulated genomes of 36.23 Morgans, split into 39 independent regions corresponding to human chromosome arms. We used a mutation rate of *μ* = 1.65 × 10^−8^ and a recombination rate of *ρ* =1 × 10^−8^ per generation per base pair. To simulate SNP data we then downsampled sequencing data to match the genotype density and allele frequency spectrum observed using Chromosome 2 of the UK Biobank data set, using 50 evenly spaced MAF bins. We generated unphased diploid individuals by randomly pairing simulated haplotypes. Ancient data was generated using a similar procedure, with two additional steps to simulate low coverage data. We first transformed the data into pseudodiploid individuals by randomly sampling one haplotype at each site. We then set each site as missing with probability m, related to a simulated coverage parameter *C* through the relationship *m* ≈ *e*^−*C*^, further described below.

### 5.2 Simulation of missingness and coverage

We simulated low coverage data by discarding a proportion *m* of the SNPs of each individual, but often report results referring to corresponding sequencing coverage parameters. To this end, we assumed a simple model where a genome of length *G* is sequenced using *N* reads of length *L*. Using this notation, the probability that a randomly selected site along the genome is not spanned by a read is:

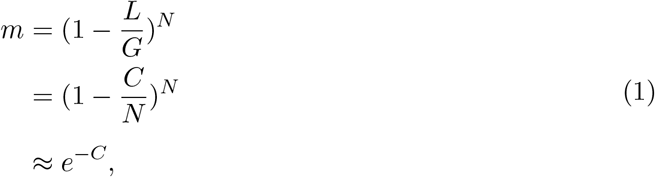

where 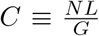 represents the coverage parameter. This relation can also be used to obtain a link between *m* and the number of reads:

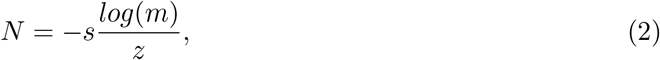

where 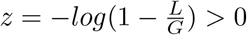 and *s* is the number of sampled individuals with missingness *m*.

### 5.3 Computation of LD

We consider a panel of *s* individuals, *M* sites and genotypes 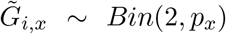 for individual *i* at site *x* with minor allele frequency *p_x_*. We first standardize the genotypes by computing 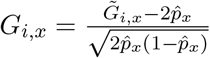, where 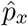 is the estimated allele frequency. The LD between two sites *x* and *y* is computed as the *R*^2^ statistic:

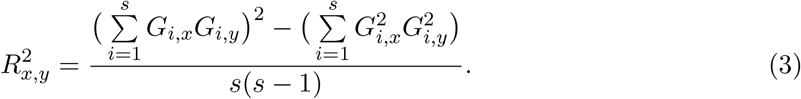

The computation of this statistic scales linearly with the number of samples 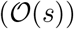. Note that this estimator is biased due to the use of 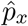 instead of the unknown allele frequency *p_x_* during the normalization step. We describe a procedure used at runtime to debias these estimates in the Supplementary Note. The LD of pseudo-diploid individuals is computed using the same approach, with 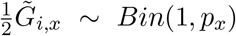.

### 5.4 Detection of IBD segments

We ran FastSMC^32^ (version 1.2) using parameters *min_m* = 0.5 (minimum cM length) and *t* = 100 (IBD time threshold). Decoding quantities were generated based on 30 samples using a European demographic history. FastSMC was run using multiple jobs, so that each job considers at most 100 haploid samples. We also used IBD segments obtained by running the HapIBD software^31^ (version 1.0), using recommended parameters for SNP-array data analysis (default parameters).

### 5.5 HapNe-IBD and HapNe-LD algorithms

We developed two algorithms to infer recent effective population size fluctuations *N_e_*(*t*) from a set of *s* samples, called HapNe-IBD and HapNe-LD. Both approaches take summary statistics {*Y_i,b_*} as input and maximize a pseudo-posterior function for *N_e_*(*t*). The input data set {*Y_i,b_*} is split into 39 genomic regions corresponding to chromosome arms indexed by *i*, using 0.5cM long bins indexed by *b*.

HapNe-IBD takes as input a list of IBD segments of length 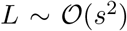. Input data {*Y_i,b_*} corresponds to the count of IBD segments in region *i* whose length lies in bin *b*. Bins start at 2cM and end at the largest detected IBD segment. We assume that each of these counts is the realization of a Poisson random variable, with demographic-dependent mean parameter *μ_b_*(*N_e_*(*t*))*L_i_* where *L_i_* is the length of the *i^th^* region (*μ_b_*(*N_e_*(*t*) is described in the Supplementary Note). To handle overdispersion, we used a quasi-likelihood approach to compute a weight parameter 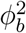 that multiplies the variance in each bin.

HapNe-LD uses average *R*^2^ statistics as input data {*Y_i,b_*}. This input is computed in 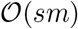, where *m* is the total number of loci. We assumed that these observations are realizations of a Normal random variable, with a distance-dependent mean parameter *μ_b_*(*N_e_*(*t*) (see Supplementary Note for a detailed description of *μ_b_*(*N_e_*(*t*))). The variance parameters 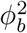 were estimated using the usual variance estimator within each bin.

Give a set of IBD or LD observations {*Y_i,b_*} for the *i^th^* genomic region and *b^th^* bin, HapNe aims to maximize *P*(*N_e_*(*t*)|{*Y_i,b_*}) under the following assumptions. First, *N_e_*(*t*) is a piece-wise exponential function from *t* = 0 to *t* = *t_max_* generations, and remains constant afterwards. In all our analyses, we used *t_max_* = 125 generations. The lengths of the time intervals are iteratively tuned so that each time interval contains the same number of expected ancestors of IBD segments (see Supplementary Note). Second, we assume that there exists a prior on the effective population size *p_N_e__*(*θ*), where *θ* represents the set of parameters defining *N_e_*(*t*). A discussion about the choice of this prior can be found in the Supplementary Note. Third, we assume that the covariance across consecutive bins can be modeled using a power likelihood 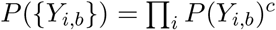. In the Supplementary Note, we show that under these assumptions the MAP estimator of *N_e_*(*t*) depends on a single hyperparameter *cσ*^2^, that we automatically tune using a heuristic model selection rule (see Supplementary Note).

Once the time intervals and the value of the regularisation parameter are fixed, HapNe assesses the uncertainty of the prediction by performing 100 bootstrap iterations. For each iteration, HapNe samples chromosome arms with replacement to create new input data, and estimates the effective population size. The 2.5th, 25th, 75th, and 97.5th percentiles are reported at each generation to obtain 50% and 95% confidence intervals.

### 5.6 Comparisons to other methods

To perform method comparisons, we simulated genotypes based on the demographic models shown in Figure 1 and used the methodology described above to compute summary statistics. We ran HapNe-IBD, HapNe-LD, IBDNe (version 23Apr20.ae9), and GONE (Jun 21, 2021 commit) using their default parameters. The simulated SNP array data did not contain enough sites to perform the SNP bootstrapping strategy used by GONE to produce confidence intervals in sequencing data. All computations were run on an Intel Skylake 2.6 GHz architecture on the Oxford Biomedical Research Computing cluster.

We reported the root mean squared log-error (RMSLE) over the first 50 generations as a measure of accuracy. If *N_e_*(*t*) and 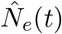 denote the true and predicted demographic models, the accuracy is defined as:

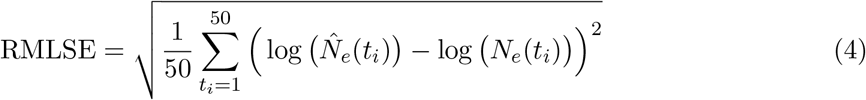

We performed 10 independent sets of simulations and computed error bars reported in each plot as 1.96 × se.

### 5.7 Filtering of high IBD and LD regions

To mitigate the impact of natural selection and structural variation, HapNe applies a filtering algorithm to exclude chromosome arms with unusual amounts of IBD sharing or LD. For LD data, parameters of a normal distribution are computed for each bin using the median and quantiles of the observed data. We used this quantile-based approach instead of moment-based estimators so that the inference is robust in the presence of the outlier regions we aim to filter out. Then, each genomic region is discarded using the following two heuristic rules. First, the deviation between the observed LD in the region and the median must be within 6 standard deviations. Second, the observed values must cross the median at least once, i.e. a region cannot have all its observations above or below the median. The IBD data is filtered using a similar approach. For each region, the mean of the Poisson distribution and the dispersion factors are computed for each bin using all others regions. The region is discarded if the sum of its squared deviance residuals is in the upper or lower *α*-quantile of the underlying χ^2^ distribution, with *α* = 10^−12^. The procedure is performed a second time, without considering the discarded regions, to prevent outliers to impact the final result.

### 5.8 LD-based admixture test

Admixture creates long-range LD between unlinked pair of sites. HapNe allows testing for the presence of admixture LD by computing cross-chromosome LD (CCLD). In the absence of CCLD, we expect the correlations between two sites *x* and *y* located on different chromosomes to be only due to finite sample sizes (see Supplementary Note):

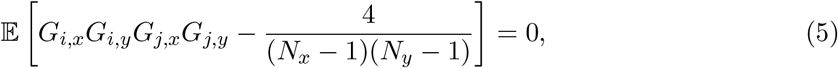

where *N_x_* and *N_y_* are the number of observed haplotypes on sites *x* and *y*, respectively. Because the LD is only computed between pairs of sites containing at least 2 overlapping observations, *N_x_* and *N_y_* are not independent variables. HapNe-LD computes the empirical mean of Eq. 5 for each pair of chromosomes and then performs a *t*-test to check for deviation from the 0-mean hypothesis. If the hypothesis is rejected, the levels of admixture LD might cause a recent collapse in the effective population size, as shown in Supplementary Figure S13.

### 5.9 Time heterogeneity in the set of analyzed samples

Most aDNA data sets contain samples originating from different time points, with an estimated date range spanning many generations when the archeological context is used to date the samples. We thus extended HapNe-LD to account for time heterogeneity and uncertainty. The user can provide a date range for each sample. This information is used by HapNe to compute the density of the ages of a randomly selected pair of individuals. This density is then used to marginalize out the age of the oldest sample and the generation gap between the two individuals under the SMC approximation, resulting in an unbiased estimator of the effective population size (see Supplementary Note).

### 5.10 Inference of demographic history in the UK Biobank

We analyzed the subset of 305,784 unrelated samples with self-reported White British ancestry, corresponding to the individuals reported in Byrcroft et al.^53^ that did not withdraw from the study and whose birth location can be assigned to a postcode in the U.K. (13,995 were removed because of this last condition). The autosomal variants were phased using Beagle 5.1^54^. We then grouped the individuals based on their self-reported birth location, labeling each of them with the first 1 or 2 letters of their corresponding postcode. We randomly picked postcodes with different sample sizes to infer population sizes. LD computations and IBD detection steps were performed using the procedure described above.

### 5.11 Inference of demographic history in the 1,000 Genomes Project

Starting with *N* = 2,504 samples from the 1,000 Genomes Project data set, we removed related individuals (up to 3*^rd^* degree) based on publicly available pedigree information. The remaining 2,460 were split according to population labels. Before running FastSMC, we downsampled the genotypes to UK BioBank as done for SNP array data, using the procedure described above. LD computations were run using all loci with MAF*>* 0.25.

### 5.12 Inference of demographic history in ancient data

We downloaded version 50.0 of the Allen Ancient DNA Resource (AADR) dataset^55^. For each analysis, we started by removing related individuals reported in the annotation files present in the dataset. For each family, the individual with the highest coverage was kept. Information about sample ages was also extracted from the annotation file and used as input for HapNe-LD. We then removed variants and individuals with low coverage (*m* > 0.8). Specific information about each population is present in the supplementary tables S1–S7.

## 6 Data availability

Genomic data sets and annotations analyzed in this study include: UK Biobank http://www.ukbiobank.ac.uk/, genetic maps ftp://1000genomes.ebi.ac.uk/vol1/ftp/technical/working/20110106_recombination_hotspots/, 1000 Genomes Project phase three https://www.internationalgenome.org/data/ and the Allen Ancient DNA Resource https://reich.hms.harvard.edu/allen-ancient-dna-resource-aadr-downloadable-genotypes-present-day-and-ancient-dna-data

## 7 Code availability

The HapNe software package is freely available at https://palamaralab.github.io/software/

## 8 Acknowledgements

We thank Juba Nait Saada and Fergus Cooper for helpful discussions and suggestions; Arjun Biddanda and Shai Carmi for comments on an early version of the manuscript; Brian Zhang and Arjun Biddanda for sharing code used for various parts of the analysis. This work was supported by the Angus McLeod Scholarship (to R.F.); NIH grant R21-HG010748-01 (to P.F.P.); and ERC Starting Grant ARGPHENO 850869 (to P.F.P.). D.R. is an investigator of the Howard Hughes Medical Institute and this work was also supported by grants from the National Institutes of Health (GM100233 and HG012287), and the John Templeton Foundation (grant 61220). This work was conducted using the UK Biobank resource (Application #43206). We thank the participants of the UK Biobank project. Computation used the Oxford Biomedical Research Computing (BMRC) facility, a joint development between the Wellcome Centre for Human Genetics and the Big Data Institute supported by Health Data Research UK and the NIHR Oxford Biomedical Research Centre. Financial support was provided by the Wellcome Trust Core Award Grant Number 203141/Z/16/Z. The views expressed are those of the author(s) and not necessarily those of the NHS, the NIHR or the Department of Health.

## Supplementary Information

### 1 Supplementary Note

#### 1.1 Derivation of the IBD and LD models

This note describes the models used to infer effective population size from IBD and LD summary statistics. We first describe a link between the effective population size and the probability that two sites are spanned by an IBD segment under the SMC’ model^1^, as well as computationally tractable approximations used in several derivations. Related work on calculations presented in this section may be found in^2–11^. We then provide details on how these models are used to perform inference based on IBD and LD summary statistics. We conclude by describing further details of the LD model related to low coverage data, time-heterogeneity, and admixture LD.

##### 1.1.1 Notation

We aim to infer the effective population size *N_e_*(*t*) based on the genotype of *s* samples consisting of *m* markers. For simplicity, we will assume that *t* is a continuous variable, with *t* = 1 corresponding to 1 generation. Note that *N_e_*(*t*) refers to haploid individuals in the population. Although *N_e_*(*t*) is the quantity of interest, we will derive several expressions in terms of its inverse 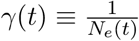, the coalescent rate, as well as the cumulative coalescent rate 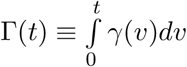.

##### 1.1.2 Survival function for a change of ancestor

Using the above notation, the distribution of the age of the most recent common ancestor (TMRCA) under the coalescent^12^ may be expressed as:

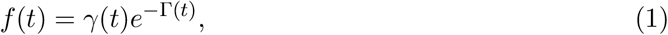

which for a constant coalescent rate takes the form of an exponential waiting time *f*(*t*) = *γe^−γt^*, leading to 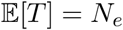.

Given the MRCA at site *x*, with TMRCA= *t*, we are interested in the genetic distance *U* at which a change of ancestor is observed. This requires a recombination event, which occurs at rate 2*t* (see e.g.^13^). When a recombination event happens, a new lineage is created at a time *V* ~ Uniform(0, *t*). This new lineage will not lead to a change of ancestor if it coalesces back to the lineage from which it branched out between *V* and *t*. We refer to this kind of coalescent event as a “healing” event and denote its probability by *ph*(*t*). To derive an expression for *ph*(*t*), we note that the coalescent rate of the new lineage is given by *f*_2_(*t*) = 2*γ*(*t*)*e*^−2Γ(*t*)^, with a factor 2 appearing because the new lineage can coalesce with either of two original ones. Healing requires the new lineage to coalesce between *v* and *t*, which happens with probability 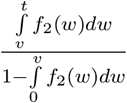. It also requires the new lineage to coalesce to the original lineage, which happens with probability 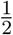. Together, these terms lead to the following expression, also derived in^7^:

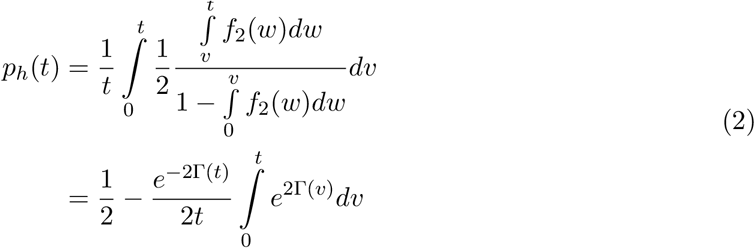

For a constant demographic history with coalescent rate *γ*, this becomes:

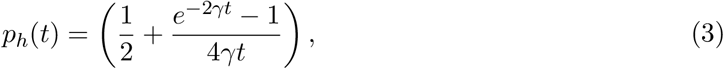

Thus, the waiting distance for a change of ancestor is exponentially distributed with rate 2*t*(1 – *ph*(*t*)) and its survival function is given by:

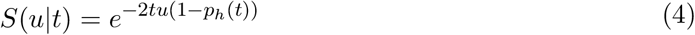

We obtain *S*(*u*) by marginalizing the TMRCA,

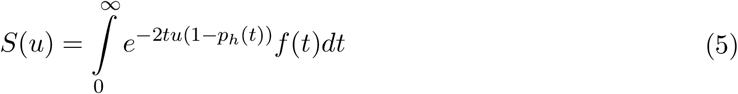

For a constant population size, this expression becomes:

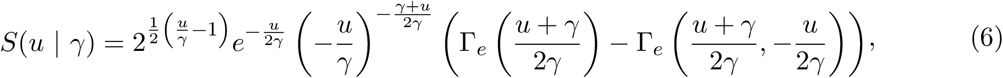

where Γ_*e*_ denotes the (incomplete) Euler gamma function 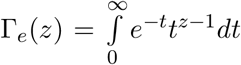 and 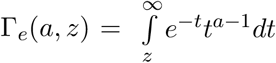. This survival function, also derived in^14^, assumes an underlying SMC’ model^1^, but does not lead to a closed-form solution when a piece-wise constant function *γ*(*t*) is utilized. To obtain a tractable expression, we introduce an approximation of the SMC’ model. Using a Taylor expansion, Eq. 4 may be written in the form:

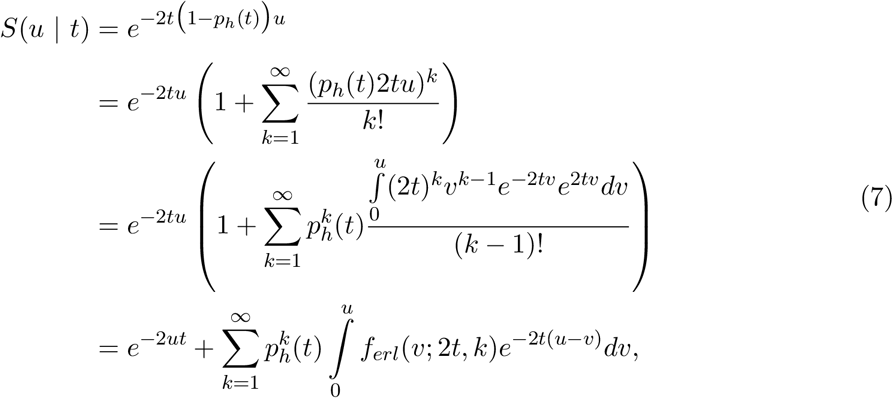

where 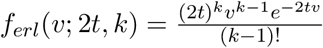 is the probability density function of the sum of *k* exponential random variables with rate 2*t*. In the last sum, *k* can be interpreted as the number of healing events observed within a distance *u*. The SMC approximation, where each recombination event leads to a change of ancestor^15^, is recovered by only considering the first term and discarding the sum:

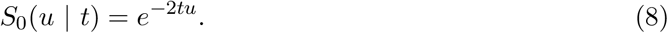

For a constant demographic history, the survival function becomes:

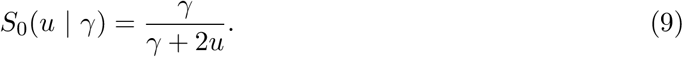

Note that this recovers the expression derived in ^16^ using a different approach. This approximation may become poor when working with small populations and short genetic distances. For example, considering *u* = 1cM and 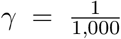 leads to a relative error 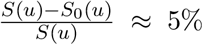. Taking into account a single recombination and healing event leads to increased accuracy (see e.g.^3^ for a related approach). Using the above formulation, this amounts to considering the first term of the sum. Under a constant demographic model, the survival function is now:

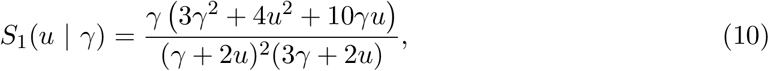

which greatly reduces the relative error compared to the SMC approximation (e.g. ~ 10 × lower using the previous example). This approach thus provides a good balance between accuracy and computational cost, as it allows multiple expressions to be computed analytically if *γ*(*t*) is approximated by a piece-wise constant function.

##### 1.1.3 IBD model

We aim to model the number of IBD segments of particular lengths shared between pairs of individuals from a population. We denote the probability density function of the length of an IBD segment by *f_seg_*(*l*|*γ*(*t*)), dropping the *γ*(*t*) term for clarity. We first consider the length of an IBD segment spanning a given site *x* along the genome. The probability density function for the length of such a segment, *f_site_*(*l*), is related to *f_seg_*(*l*) through the following relation^2^:

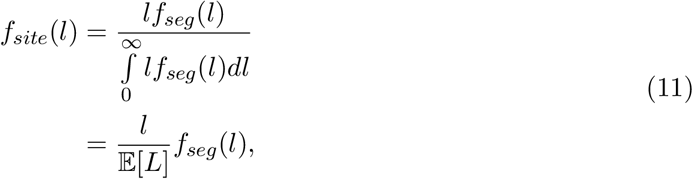

where 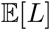 represents the expected length of a randomly selected IBD segment. The TMRCA of the two haplotypes at site *x* is distributed according to *f*(*t*). Conditioned on a TMRCA *t*, the length of the IBD segments spanning *x* is the sum of the distances to the next change of ancestor on either side of the site. By allowing at most one healing event within the IBD segment as described above, the density takes the form:

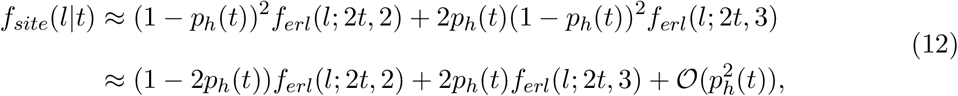

where the first term accounts for the case of no healing events and the second term allows for one recombination event. Marginalizing *t*, we obtain:

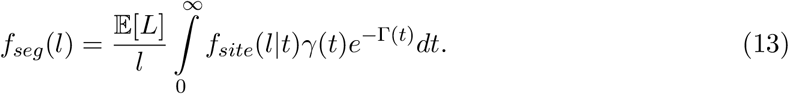

For a constant demographic history, this becomes:

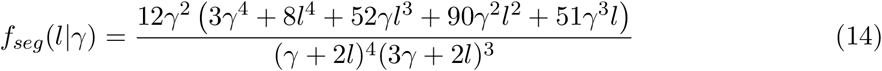

Neglecting the probability of healing leads to the SMC approximation for a constant demographic history:

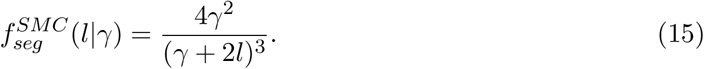

Conditioned on the total number of IBD segments *N_s_* shared in a region, the expected count of IBD segments within a length bin delimited by *u_i_* and *u*_*i*+1_ is 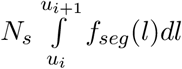. Furthermore, 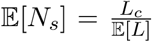, with *L_c_* denoting the genomic length of the current region. Thus, the expected value of the number of segments within the *i^th^* bin *Y_i_* is given by:

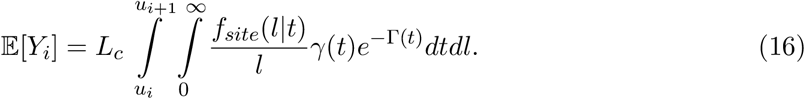

Note that we neglect issues due to finite size chromosomes, which we found to have a negligible effect. For a constant demographic history, this quantity becomes:

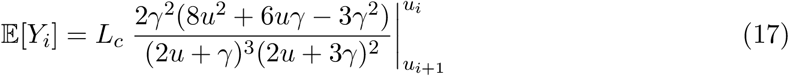

Eq. 16 provides the first moment of the distribution of *Y_i_*. Note that the approximation introduced in Eq. 10 allows to compute this expression analytically when the demographic model *γ*(*t*) is a piece-wise constant function. Previous expressions derived under the full SMC’, on the other hand, required the use of special functions or numerical integration^7^.

Poisson distributions provide a natural way of describing “count data” such as *Y_i_*. However, when using the Poisson model, we encountered bin-dependent overdispersion, particularly for smaller bins, where IBD segments originate from older coalescence events that likely involve multiple samples. We thus used a quasi-likelihood approach^17^, adding a dispersion parameter *ϕ_i_*:

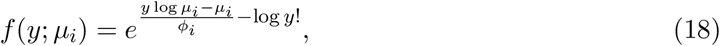

where 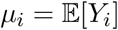 and the Poisson mass function is recovered for *ϕ_i_* = 1. The dispersion parameters *ϕ_i_* are set so that the variance of the deviance residuals is 1.

##### 1.1.4 LD model

Rather than relying on the direct observation of IBD data, HapNe-LD leverages long-range correlations that are induced by shared segments, which may be detected using unphased data. To describe the LD model used by HapNe, we begin by noting that alleles found at high frequency in a sample are typically older than ancestors transmitting large IBD segments (also see Section 1.2.1 for calculations related to the age of IBD segments). This implies that high frequency mutations found on long IBD segments are also likely to be carried by the shared ancestor transmitting the segment. We restrict our analysis to sites with MAF > 0.25. Given one such high frequency site *x*, we assume that the haplotypes of two individuals *i* and *j* spanned by a large (> 0.5 cM) IBD segment satisfy

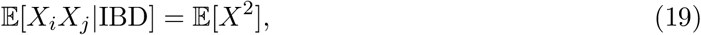

and that the same haplotypes will be independent if not spanned by an IBD segment, i.e.

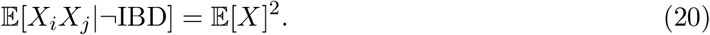

The presence of IBD segments therefore leads to correlation in the observed genotypes, which HapNe-LD aims to leverage for the inference of effective population size variation. The input for HapNe-LD is a set of unphased genotypes 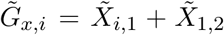, where *i* ∈ {1, …, *s*} denote individuals in the panel, and *x* ∈ {1, …, *M*} denote sites. 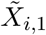 and 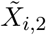 represent the (hidden) haplotypes of sample *i* at site *x*, with 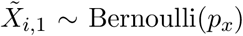 where *p_x_* is the population’s allele frequency at site *x*. For simplicity, we consider standardized input data:

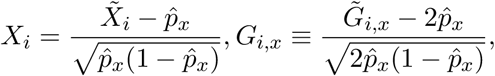

where 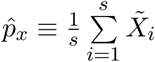 is the estimator of the allele frequency at site *x*, which is assumed to remain constant in the recent past.

HapNe-LD starts by computing the LD for different bins *b*. Unless otherwise specified, these bins are 0.5cM long and range from 0.5 to 10cM. For every bin *b*, we compute 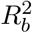 as the average of all 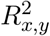 values estimated for pairs of sites (*x, y*) whose distance is within bin *b*:

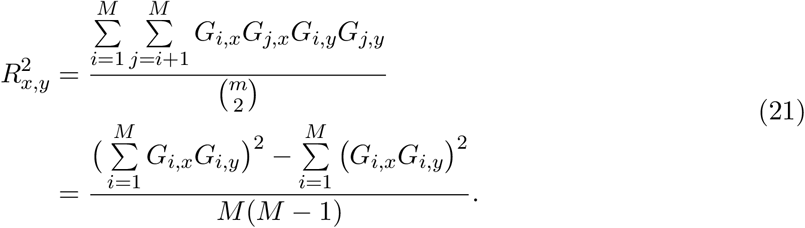

Note that this requires 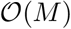 computation.

We now aim to relate these correlation statistics to the effective population size. The first moment of 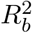 is given by:

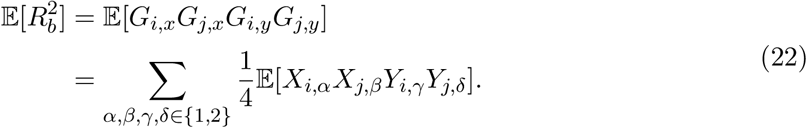

We can group the 16 terms of the sum into different categories, according to the number of distinct haplotypes involved in each of these terms. In particular, the 4 terms where *α* = *γ* and *β* = *δ* involve two distinct haplotypes, i.e. haplotype *α* for individual *i* and *β* for individual *j*. For these 4 terms, we can use equations 10, 19, and 20 to write:

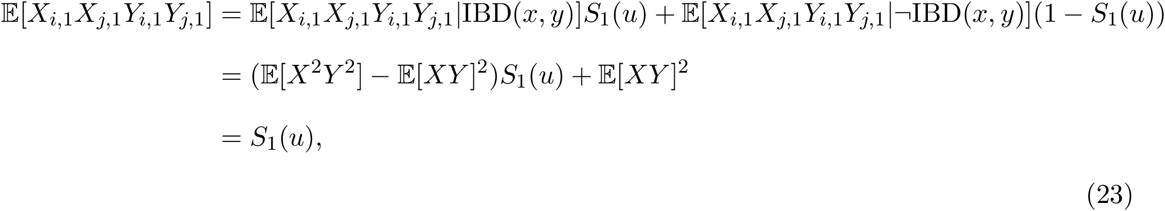

where *u* denotes the distance between the two sites *x* and *y*. Note that we neglect issues due to finite sample sizes and admixture LD, which are addressed later. With this assumption, we have 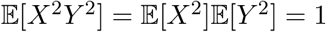 and 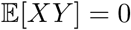.

The 12 other terms of the sum of Eq. 22 involve either 3 or 4 haplotypes. For example, a term with *α* ≠ *γ* and *β* = *δ* involves both haplotypes for individual *i* and haplotype *β* for individual j. In these cases, correlations induced by IBD require at least two pairs of haplotypes to be shared IBD, leading to 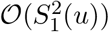 contributions, which we neglect.

Together, these expressions enable obtaining the first moment of 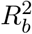. If bin *b* is delimited by *u_i_* and *u_j_*, we have:

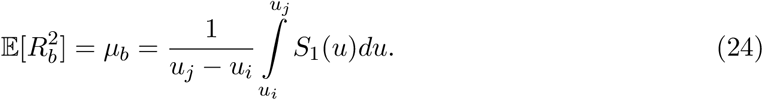

To complete the model, we assume that

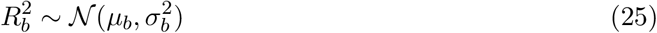

and estimate 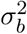 using 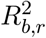 estimates obtained across chromosome arms.

##### 1.1.5 Correcting for finite sample size

Working with finite sample sizes induces correlations in the data which, if not accounted for, lead to bias in the inferred effective population size. These correlations arise as a result of the use of an empirical allele frequency 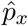 instead of the unknown *p_x_*. As a first step to debias the estimator of *R*^2^, we consider the ratio of the expected values as an approximation to the expected value of the ratio, which has been shown to be a good approximation for common alleles^18^:

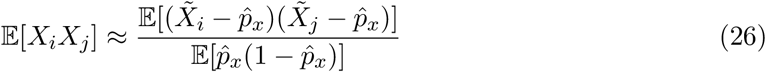

If *s_x_* haplotypes are observed at site *x*, the numerator becomes:

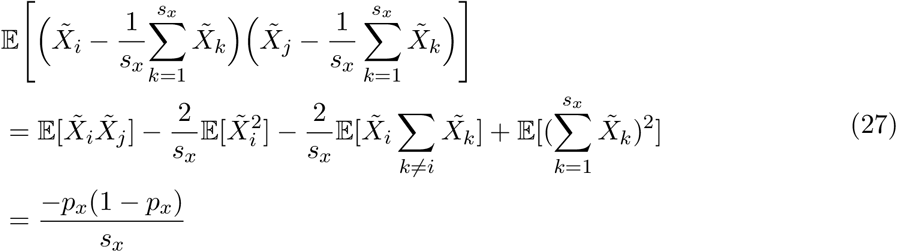

Similarly, the denominator is given by:

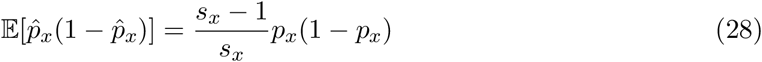

It follows that:

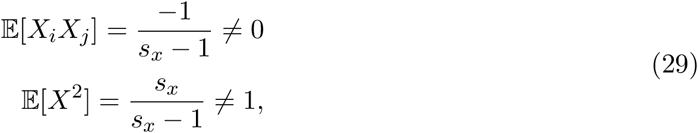

When working with low coverage data, *s_x_* becomes a random quantity, *S_x_*. Because computing LD between *x* and *y* requires that at least two individuals are sequenced at both sites, *S_x_* and *S_y_* are not independent for the (*x, y*) pairs considered when computing LD. We therefore average realizations of 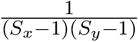 over pairs of sites (*x, y*) to compute an estimate 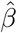 for the following quantity in Eq. 23:

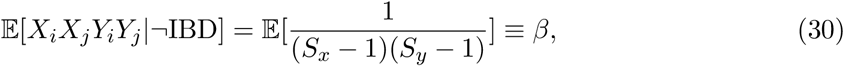

which is also relevant for the detection of admixture LD, as discussed later. We use the same pairs (*x, y*) to similarly obtain an estimate 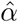 for the quantity

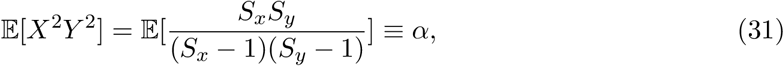

and use these terms to obtain a corrected estimate for 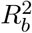

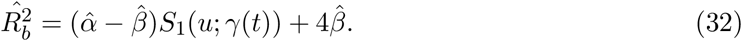

Note that the factor 4 is due to the 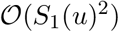 terms in Eq. 22 that also cause finite-sample size correlations.

##### 1.1.6 Correcting for time heterogeneity

Ancient DNA samples in a data set often originate from different time points. Due to the uncertainty in obtaining precise time estimates, their origins are often reported as a time range. Time heterogeneity across the set of analyzed samples causes a reduction in LD, due to the effects of recombination on the underlying haplotypes. If not modeled, this leads to an upwards bias in the estimated effective population size. HapNe-LD implements a correction to prevent these biases using the reported sample ages, which are obtained via radio-carbon dating or using the archeological context.

Consider two individuals *i* and *j* sampled at times *T_i_* and *T_j_*. Assume, without loss of generality, that *T_i_* > *T_j_* and define Δ*T* ≡ *T_i_* – *T_j_* > 0. Following the lineage of individual *j* at a site *x*, we denote by *k* the ancestor living at generation *T_i_*. The LD between individuals *i* and *k*, both of them living at generation *T_i_*, can be computed using Eq. 7 by replacing *γ*(*t*) with *γ_o_*(*t*) = *γ_o_*(*t* + *T_i_*). The LD between individuals *i* and *j* is obtained by multiplying the LD between individuals *i* and *k* by the probability that the haplotype is not broken by a recombination event when transmitted from *k* to *j*, which decays exponentially with rate Δ*T*. Under the SMC approximation, this probability is given by *e*^−Δ*Tu*^. In practice, *T_i_* and *T_j_* are not known exactly but provided as a range. If the density functions of *T_i_* and *T_j_* are available, both times can be marginalized in the above calculations of LD. HapNe supports used-provided time intervals for each sample and assumes that the true time is uniformly distributed within these intervals.

##### 1.1.7 Admixture LD

Admixture causes correlation due to differences in allele frequencies across diverged populations. This correlation, often referred to as admixture LD, may lead to biases in the inferred demographic models. We use Eq. 30 to detect the presence of admixture LD and partially correct for it. For each pair of distinct chromosomes *i* and *j*, we compute the average difference between both sides of Eq. 30 and use a two-sided *t*-test to verify that they do not significantly deviate from 0. To mitigate the effects of admixture LD, we estimate 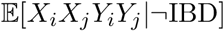 by averaging realizations of *X_i_X_j_Y_i_Y_j_* for loci located on different chromosomes, and used this value as an estimate of *β* in Eq. 32. Note that, because all pairs of chromosomes are used to compute the *t*-test, the samples are not strictly independent, making this approach slightly conservative. An alternative approach consists in only considering disjunct pairs of chromosomes, which however leads to higher variance in the estimates for *β*.

##### 1.1.8 Effective population size in IM and ICF models

We used the backward-in-time Markov chain introduced in^19^ to convert coalescence rates for the IM and ICF multi-population models into effective sizes for an equivalent single-population model. In particular, given a demographic model involving multiple populations, we used a Markov chain to compute the probability that two lineages coalesce at generation *t*, conditioned on not having coalesced up to generation *t* – 1, and took the inverse of this probability to be the effective population size for an equivalent single-population model.

#### 1.2 Additional details on the inference procedure

We provide additional details on the use of quantiles of the IBD segment age distribution to discretize the time intervals and on the regularized loss function minimized by HapNe to infer *N_e_*(*t*).

##### 1.2.1 Parameterization of *N_e_*(*t*)

HapNe aims to infer the demographic model given by *N_e_*(*t*). We parameterize this function by assuming it to be piece-wise exponential, with parameters described by a vector, *θ*. More in detail, we divide the time axis into *M* consecutive intervals and for each interval *i* assume that *N_e_*(*t*) varies according to a constant exponential rate λ_*i*_. We set λ_*M*_ = 0, implying that the population size remains constant from the last predicted time to infinity. *N_e_*(*t*) is thus fully determined by a set of *M* values 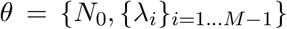. Time intervals are automatically selected so that each of them contains the same expected number of IBD segments (as also done in e.g.^20^). Let *f_age_*(*t*|*l* > *u_min_*) denote the probability density function of the age of IBD segments whose length satisfies *l* > *u_min_*. We define time intervals so that they coincide with quantiles of this density, which we compute using

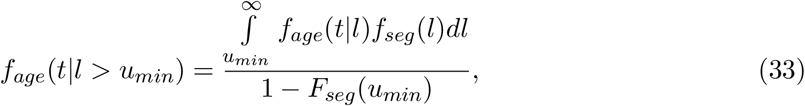

where *f_seg_*(*u*) in defined in Eq. 13 and 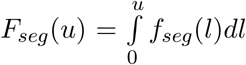. To derive *f_age_*(*t*|*l*), we note that it represents the TMRCA of a randomly selected site spanned by an IBD segment of length *l*. Using Bayes’ rule and the SMC approximation,

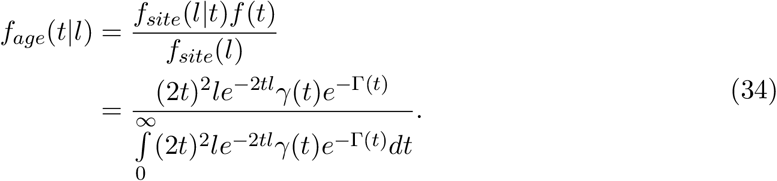

For a constant coalescent rate *γ*, this becomes

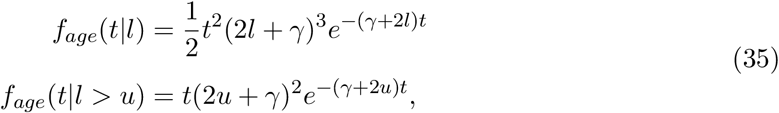

i.e. an Erlang-3 and Erlang-2 distribution, respectively (also see^6,9^). Because time intervals depend on *N_e_*(*t*), HapNe iteratively tunes them at each iteration using the current population size estimates.

Note that a slightly more accurate closed-form solution under a constant population size can be obtained by allowing a single recombination event to heal, replacing *f_site_* in Eq. 34 with the expression of Eq. 12, leading to:

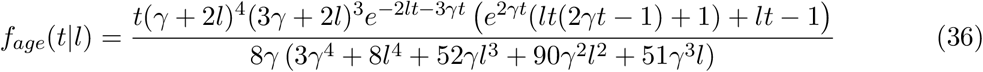

##### 1.2.2 Loss function

We aim to find the best set of parameters *θ* based on correlated observations *Y* = {*y_r,b_*}, where *y_r,b_* represents LD or IBD summary statistics computed for the *b^th^* bin of the *r^th^* independent genomic region. Due to the presence of correlations in the data, rather than using standard likelihood calculations we work with the approximated power likelihood

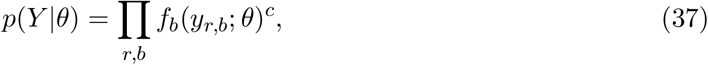

where 0 ≤ *c* ≤ 1 is a hyperparameter and *f_b_* is the probability mass or density function derived in equations 18 and 25. Minimizing Eq. 37 for *θ* is an ill-defined problem, for which small changes in the input data might lead to significant changes in the inferred parameter 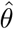 (also see e.g.^4^). To improve convergence and restrict the parameter space we thus impose the following prior on the {λ} coefficients of the piece-wise exponential function *N_e_*(*t*):

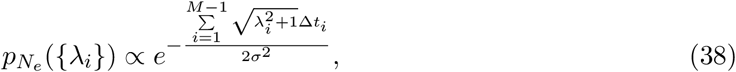

where Δ*t_i_* denotes the length of the *i^th^* time interval and *λ_i_* the growth rate in the same interval. Because the numerator corresponds to the length of log *N_e_*(*t*) between *t* = 0 and the last predicted time, this choice of prior favors trajectories with reduced fluctuations.

Combining these expressions leads to the following posterior:

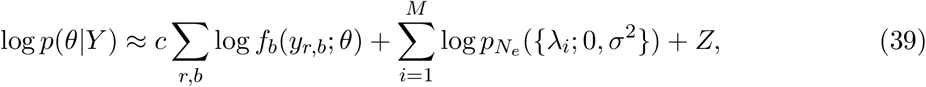

where *Z* is a normalizing constant.

We aim to find the MAP of *θ*:

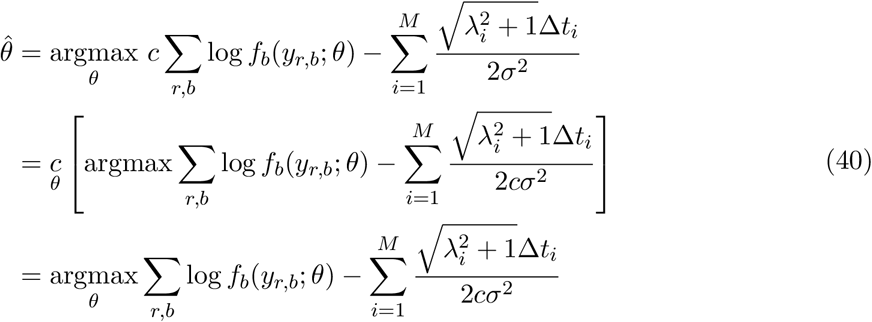

This requires tuning a single hyperparameter *κ* = *cσ*^2^, using the approach described in the next section.

##### 1.2.3 Numerical optimization

We used SciPy’s implementation of the L-BFGS-B optimiser^21^ to minimize Eq. 40. Each minimization step is run 5 times using different starting points. The solution yielding the smallest loss is kept.

#### 1.3 Model selection

HapNe performs a grid-search over different values of the hyperparameter *κ*, ranging from a strong regularization *κ*_0_ = 10^−5^ to an almost unregularized model with parameter *κ_max_* = 100. For each of these parameters, HapNe infers the MAP 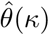 by optimizing Eq. 40, as well as the associated pseudo-likelihood 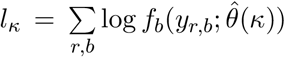. HapNe then computes the “pseudo-deviance” 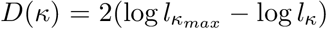. The smallest value of *κ* satisfying *D*(*κ*) < *τ* is selected as the best hyperparameter. Since the parameter *c* handling correlations between bins is neglected when computing the “pseudo-deviance”, we cannot use asymptotic theories about the distribution of *D* to fix the value of *τ* in a principled way. Instead, we fixed the thresholds *τ* for both HapNe-LD and HapNe-IBD by training them using three sets of simulations that used different demographic models than the ones presented in this work.

#### 1.4 Supplementary Figures

**Figure S1:**
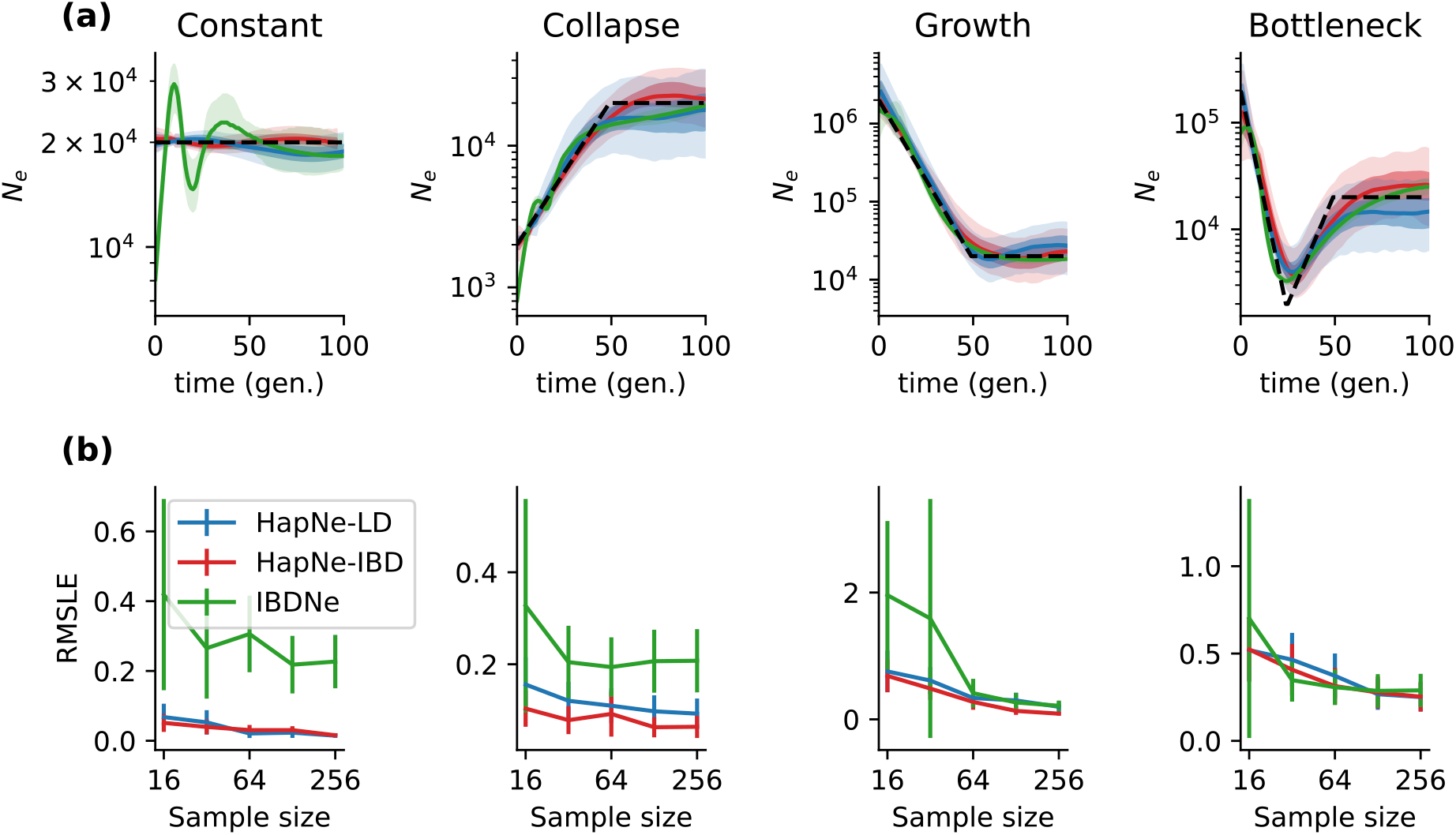
Accuracy of HapNe-IBD and IBDNe using ground truth IBD sharing information, and HapNe-LD using inferred LD. (a) Simulated demographic models (dotted black lines), predictions based on ground truth IBD sharing for both HapNe-IBD (red) and IBDNe (green), and HapNe-LD results based on simulated SNP-array data (blue). (b) Error as a function of sample size for corresponding demographic models in (a), measured as the RMSLE over the first 50 generations (see Methods). HapNe-IBD and IBDNe were run using ground truth IBD sharing information. Error bars correspond to 1.96 × *SE* computed using 10 independent simulations.

**Figure S2:**
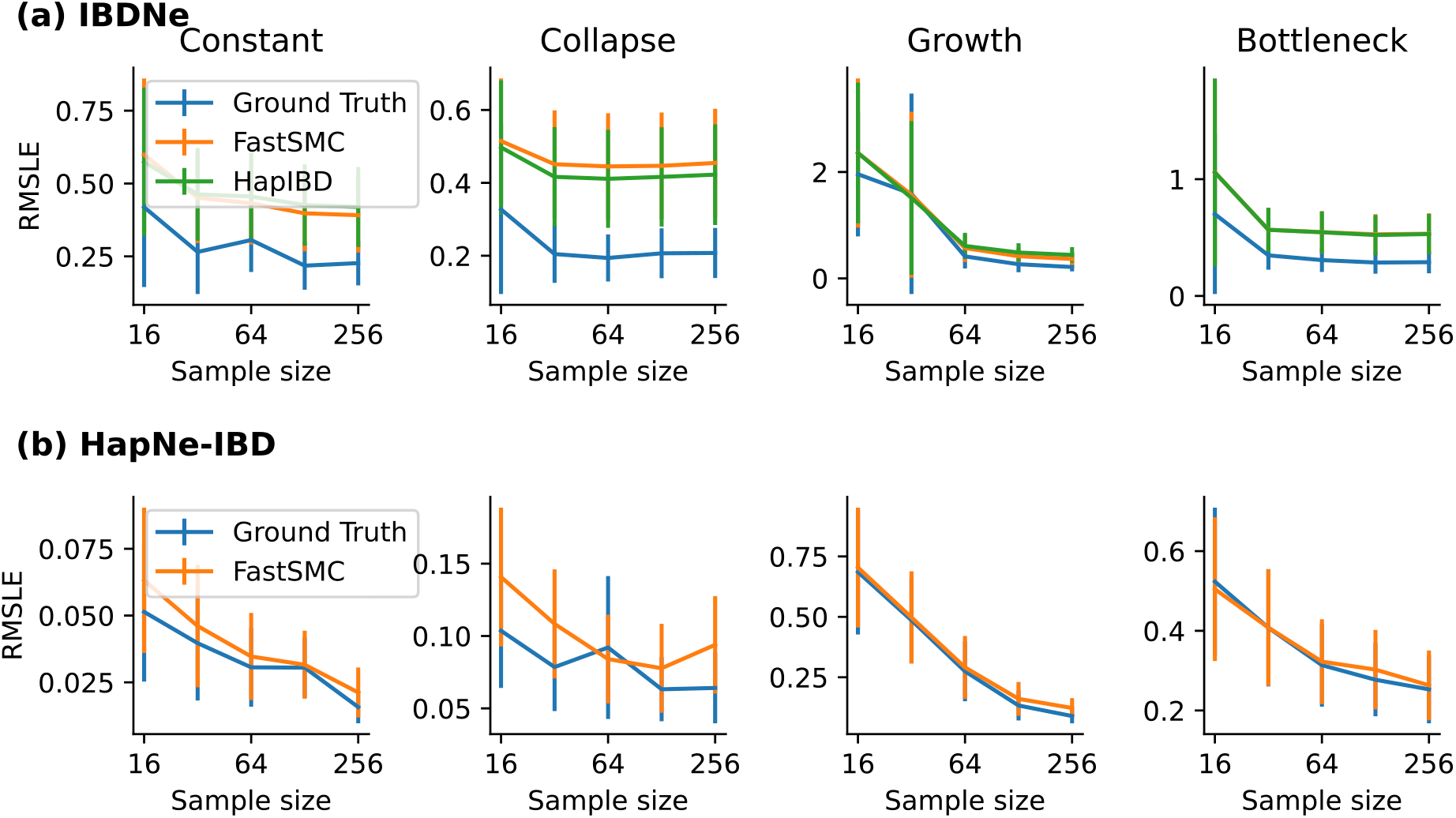
Impact of IBD detection on the accuracy of IBDNe and HapNe-IBD. (a) RMSLE as a function of sample size for IBDNe and (b) HapNe-IBD using different sources of IBD sharing. Ground Truth refers to the IBD segments obtained from the ARGON simulator, FastSMC and HapIBD were applied as described in the Methods section. Error bars correspond to 1.96 × *SE* computed using 10 independent simulations.

**Figure S3:**
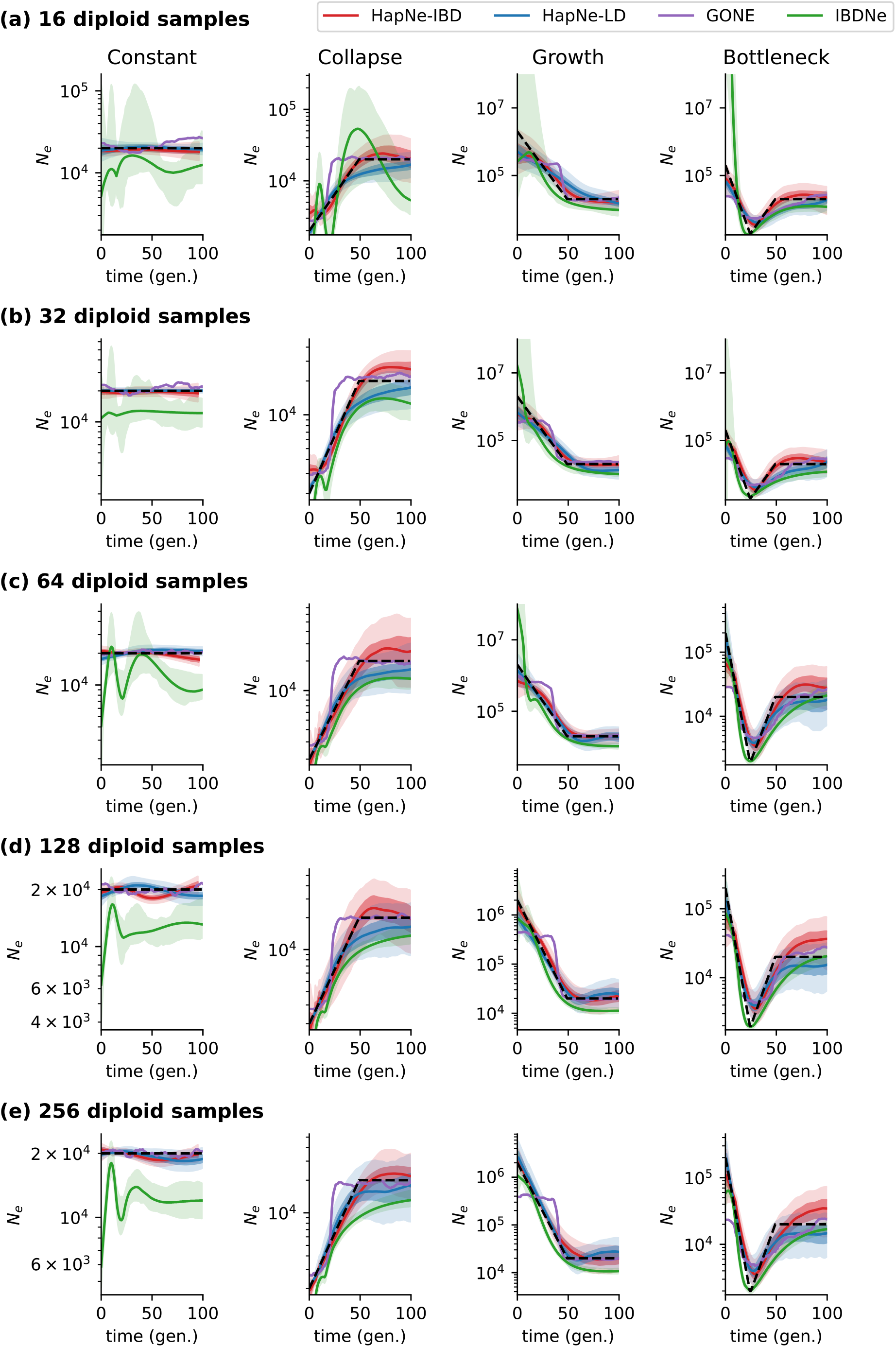
Effect of sample size variation (rows) across several demographic models (columns). HapNe-IBD was run using IBD segments detected by FastSMC and IBDNe using segments detected by HapIBD. LD methods were run using their standard pipeline. The y-axis is truncated for readability in simulations that resulted in very large vaues.

**Figure S4:**
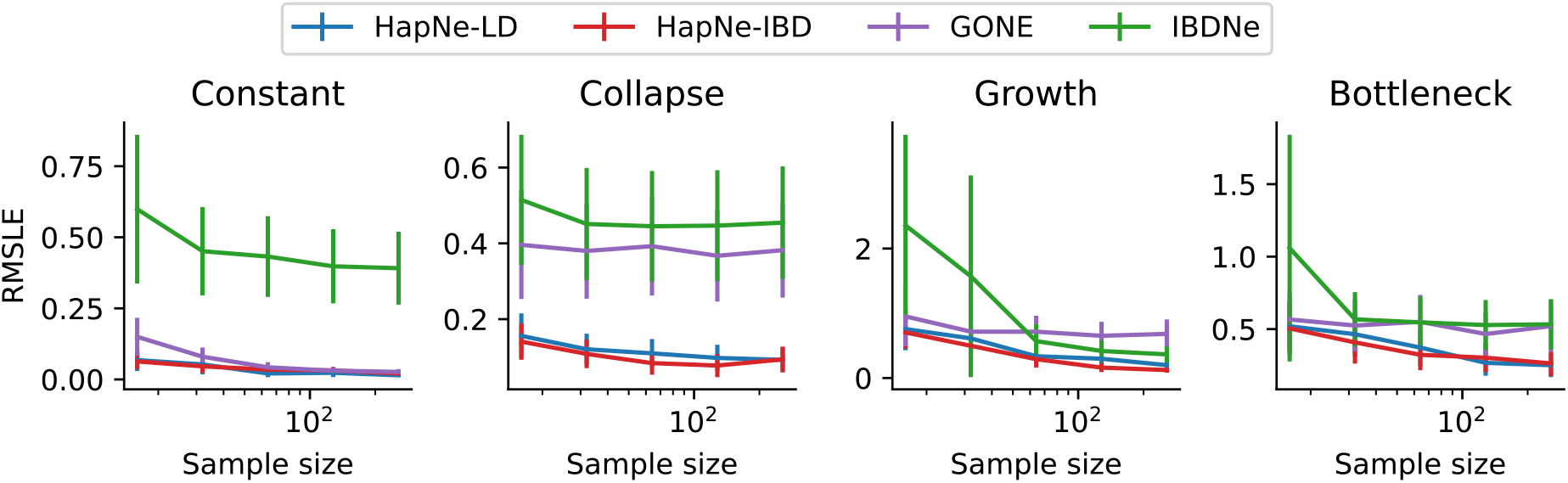
Inference accuracy as a function of sample size. Accuracy was measured using RMSLE over the first 50 generations for each simulated demographic history and sample size (see Methods). IBD segments for HapNe-IBD and IBDNe were computed using FastSMC and HapIBD, respectively. Error bars correspond to 1.96 × *SE* computed using 10 independent simulations.

**Figure S5:**
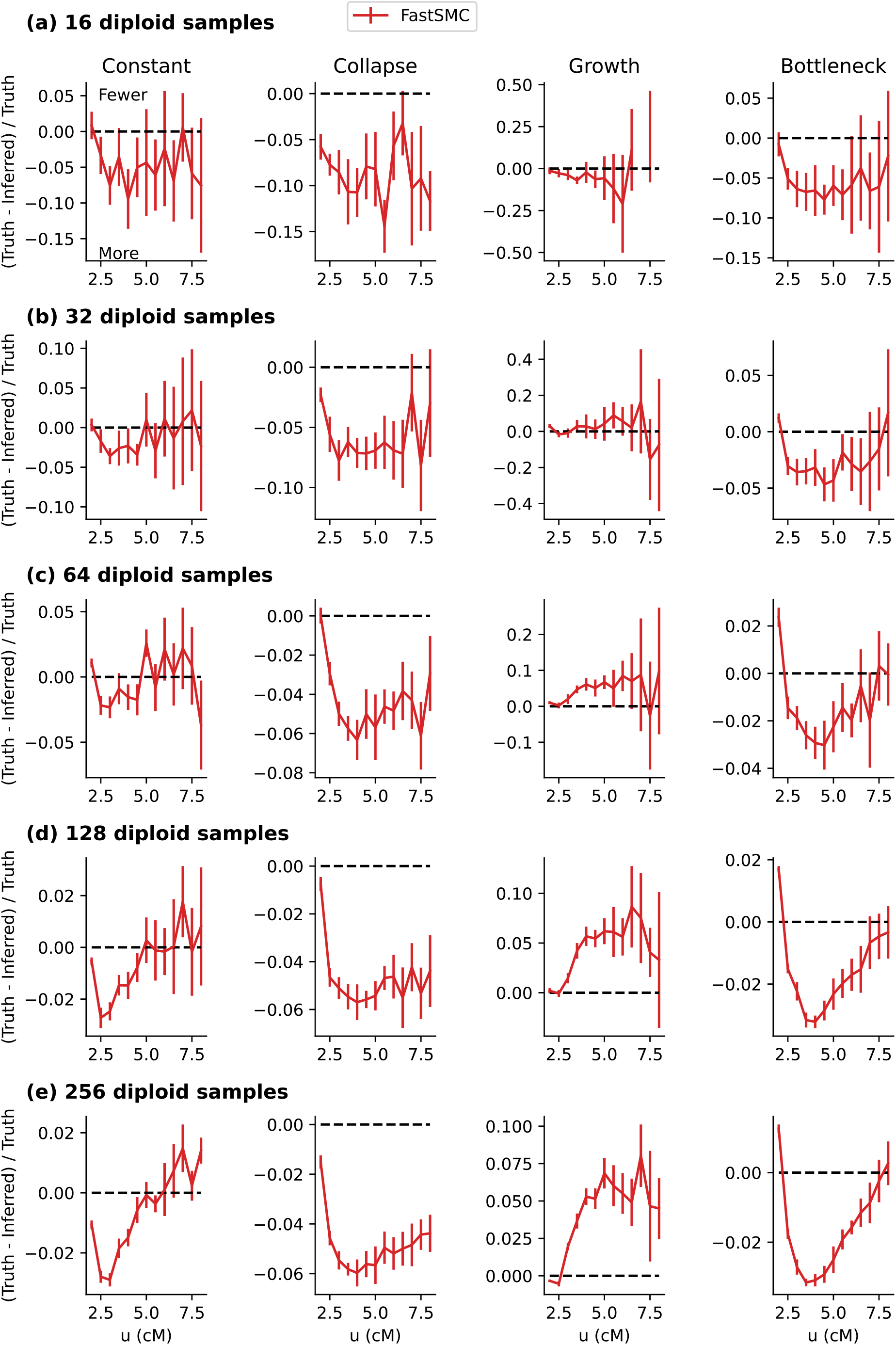
Relative error in IBD detection. We computed the relative difference between the true and inferred number of IBD segments for different sample sizes (rows) and demographic models (columns) for FastSMC. Positive/negative values indicate a depletion/excess of detected segments.

**Figure S6:**
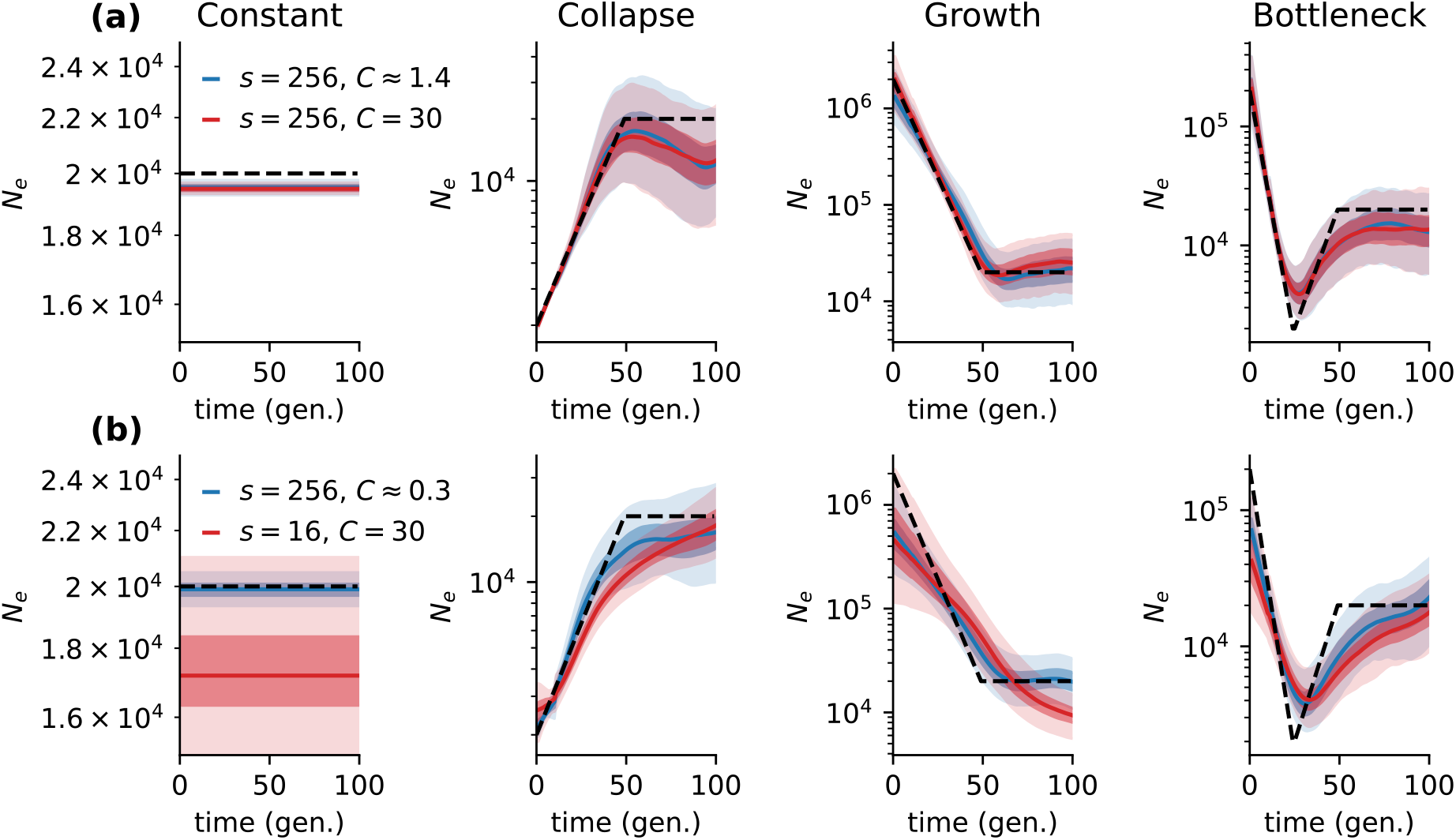
Effect of coverage and sample size. (a) Output of HapNe-LD on simulated aDNA for 256 individuals, with *m* = 0 (*C* ≈ 30) and *m* = 0.25 (*C* ≈ 1.4). (b) Output of HapNe-LD on simulated aDNA for 16 individuals with *m* = 0 (*C* ≈ 30) and 256 individuals with *m* = 0.75 (*C* ≈ 0.3).

**Figure S7:**
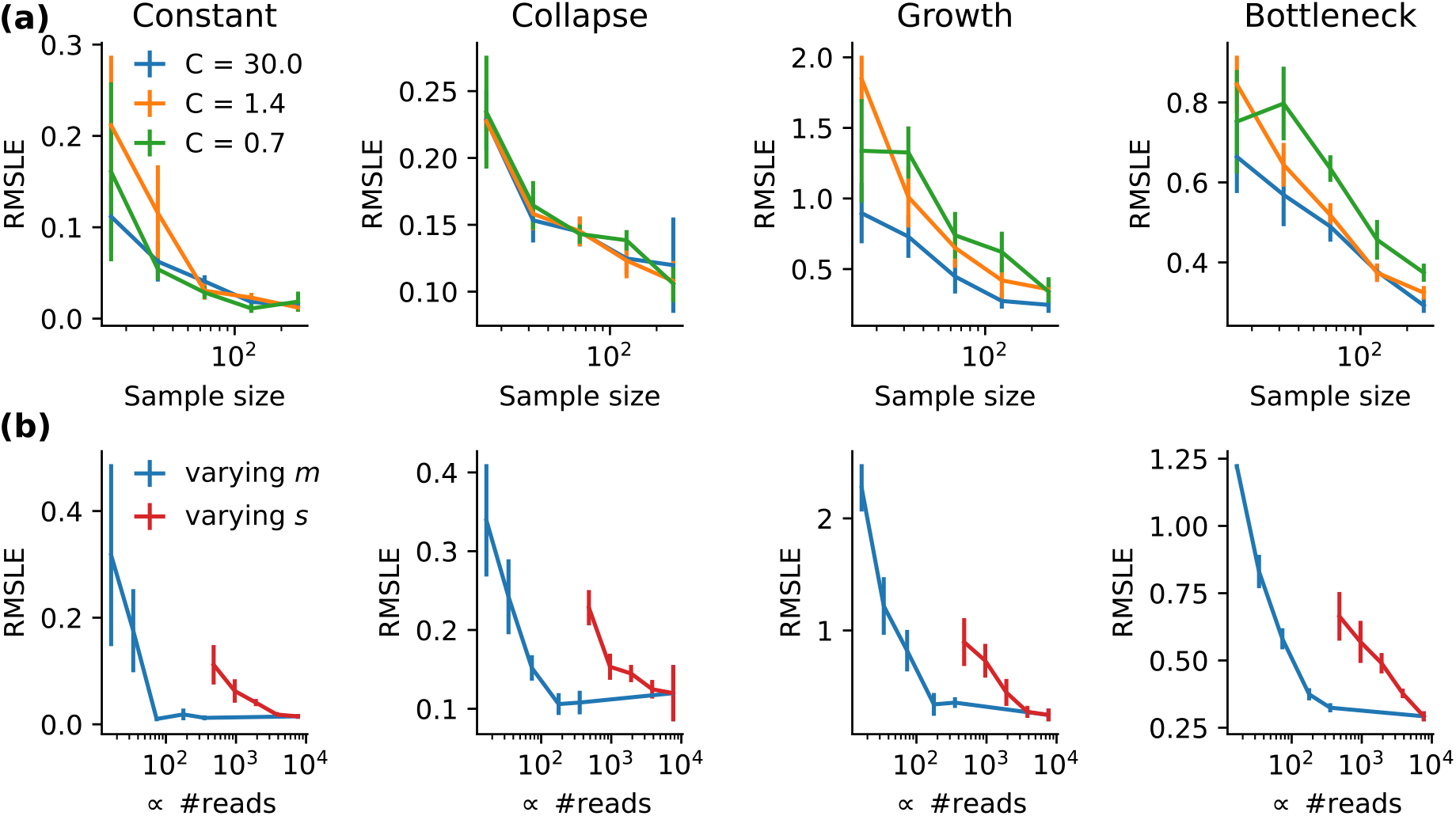
Accuracy of HapNe-LD as a function of sample size and coverage. (a) RMSLE for HapNe-LD as a function of sample size for three different levels of coverage (line color) and different demographic models (column). The different levels of coverage, 30×, 1.4× and 0.7×, approximately correspond to *m* = 0, *m* = 0.25 and *m* = 0.5, respectively (see Methods). (b) Comparison of the RMSLE while keeping the number of samples constant (*s* = 256) and decreasing coverage (blue line), compared to the RMSLE obtained while keeping the coverage constant at 30×, while decreasing the sample size.

**Figure S8:**
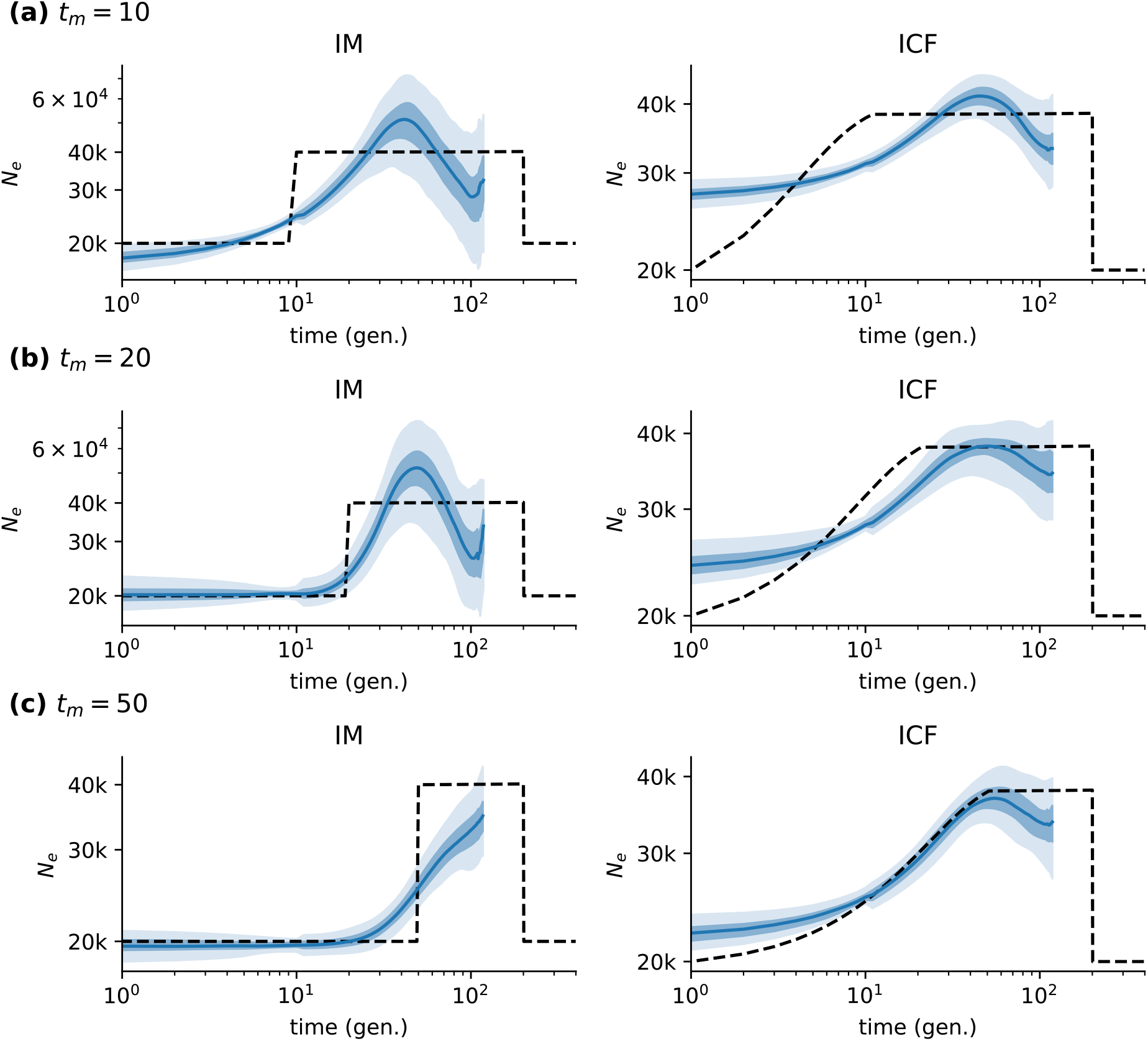
Inference based on demographic models involving multiple populations. (a-c) Results for the IM and ICF models for different values of *t_m_* (see Methods).

**Figure S9:**
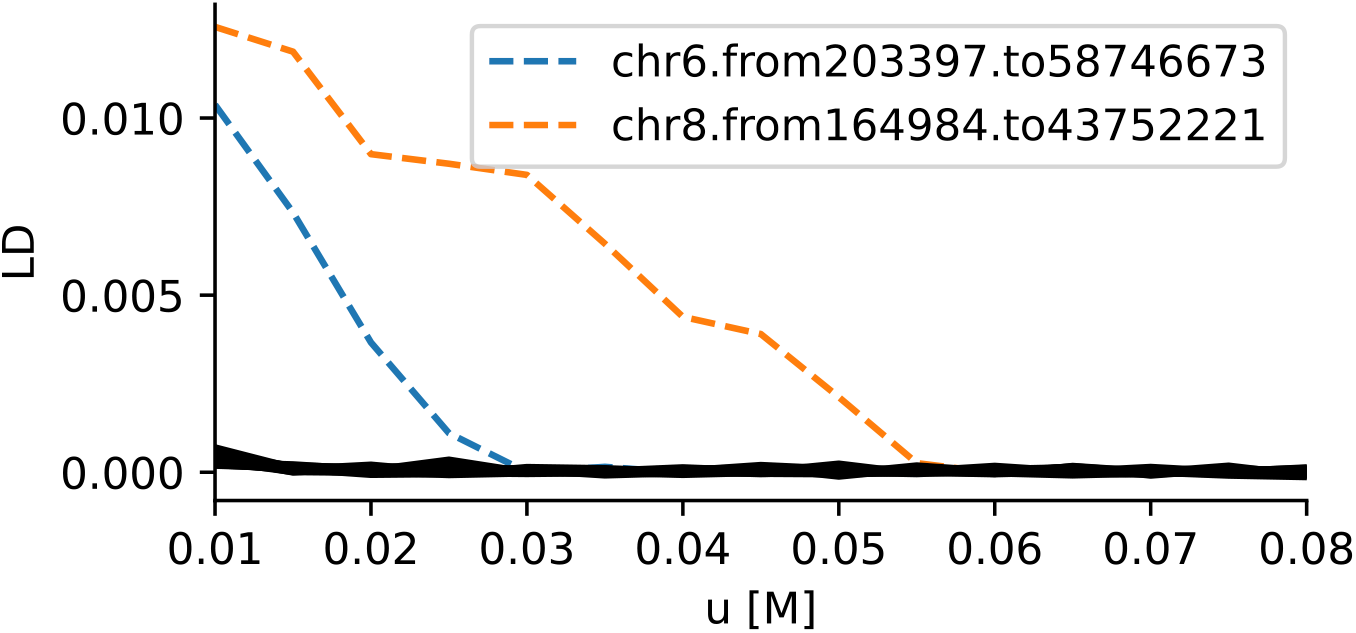
Filtering of high LD regions. The LD at different distances *u* (in Morgans, M) was computed by randomly selecting individuals from the UK Biobank. Unusually elevated LD was observed in the HLA region on Chromosome 6 (blue line) and on Chromosome 8 (orange line), corresponding to a known large inversion polymorphism.

**Figure S10:**
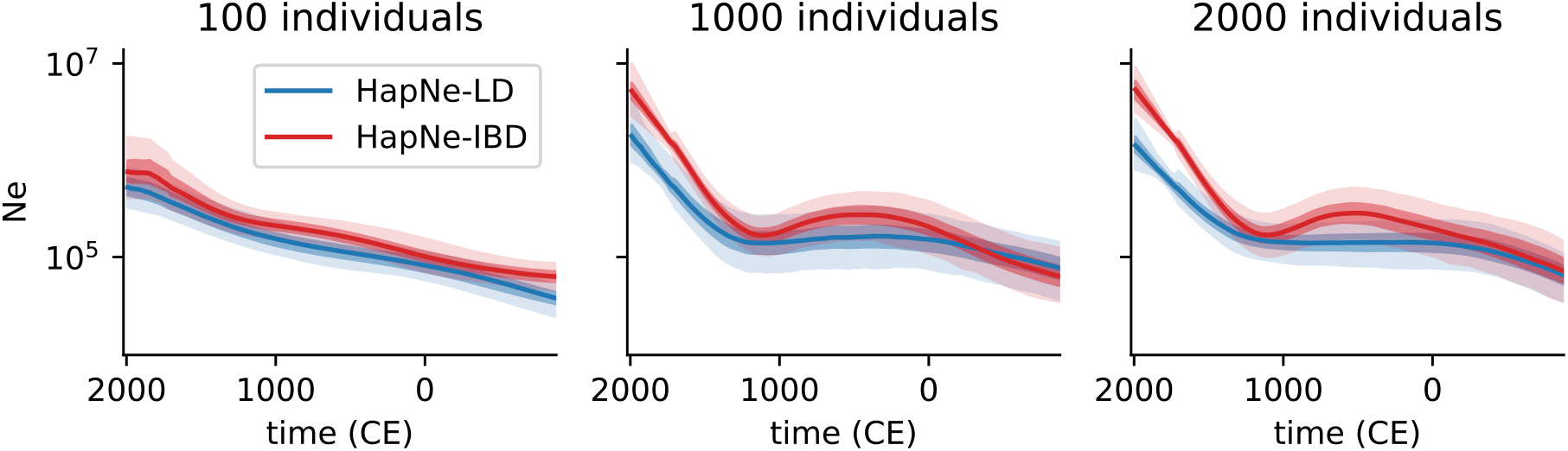
Downsampling analysis for the Glasgow postcode in the UK Biobank. Effective population size inferred using unrelated individuals with self-reported white British ancestry whose birth location is in the Glasgow (G) postcode area. The numbers above each plot correspond to the sample size used in each analysis.

**Figure S11:**
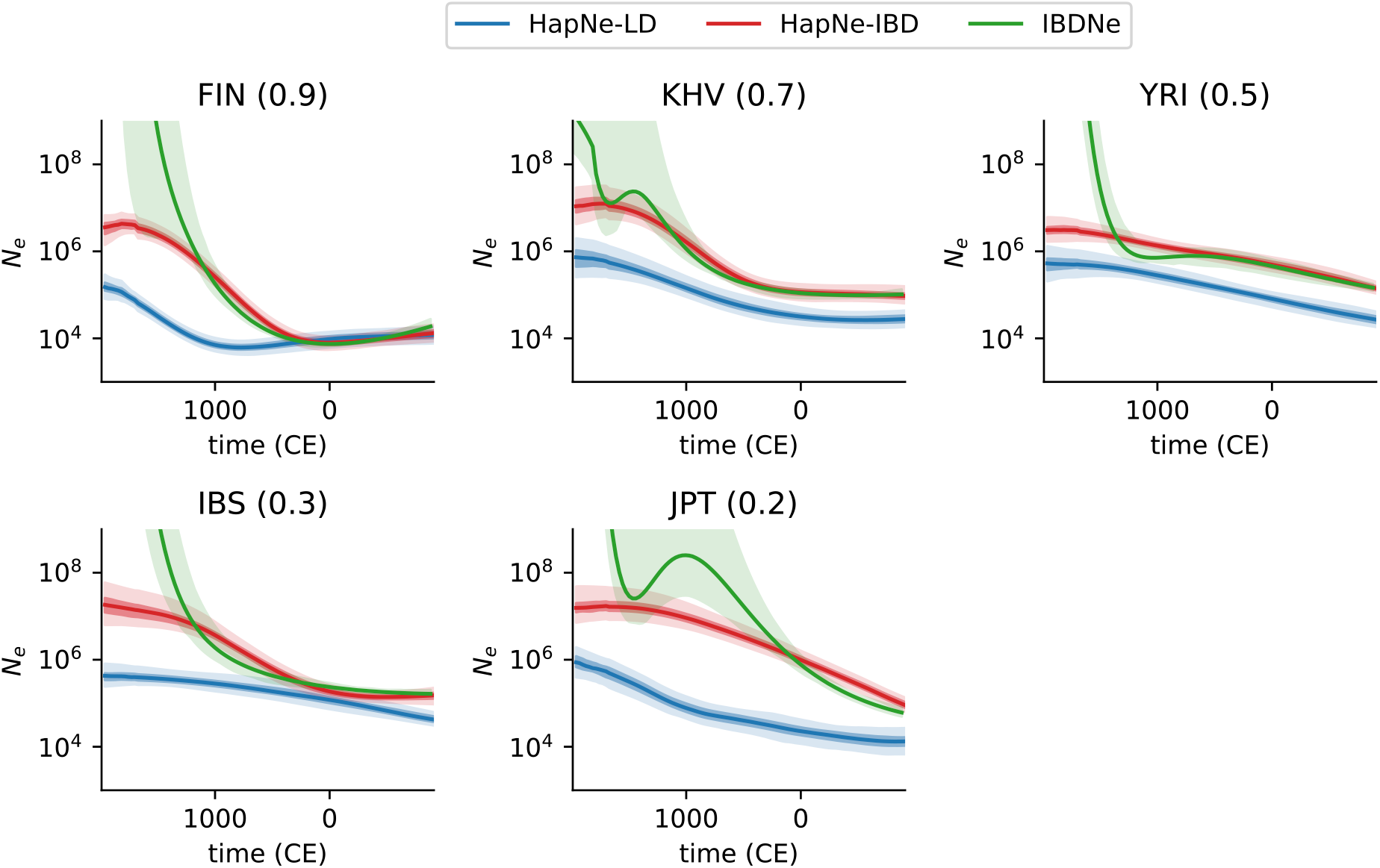
Inferred demographic models for 1,000 Genomes Project populations where no significant admixture LD was detected. Results for populations for which the admixture LD test was not significant (*p* > 0.05). Numbers in parentheses correspond to −log_10_(*p*). IBD segments for IBDNe and HapNe-IBD were computed using FastSMC.

**Figure S12:**
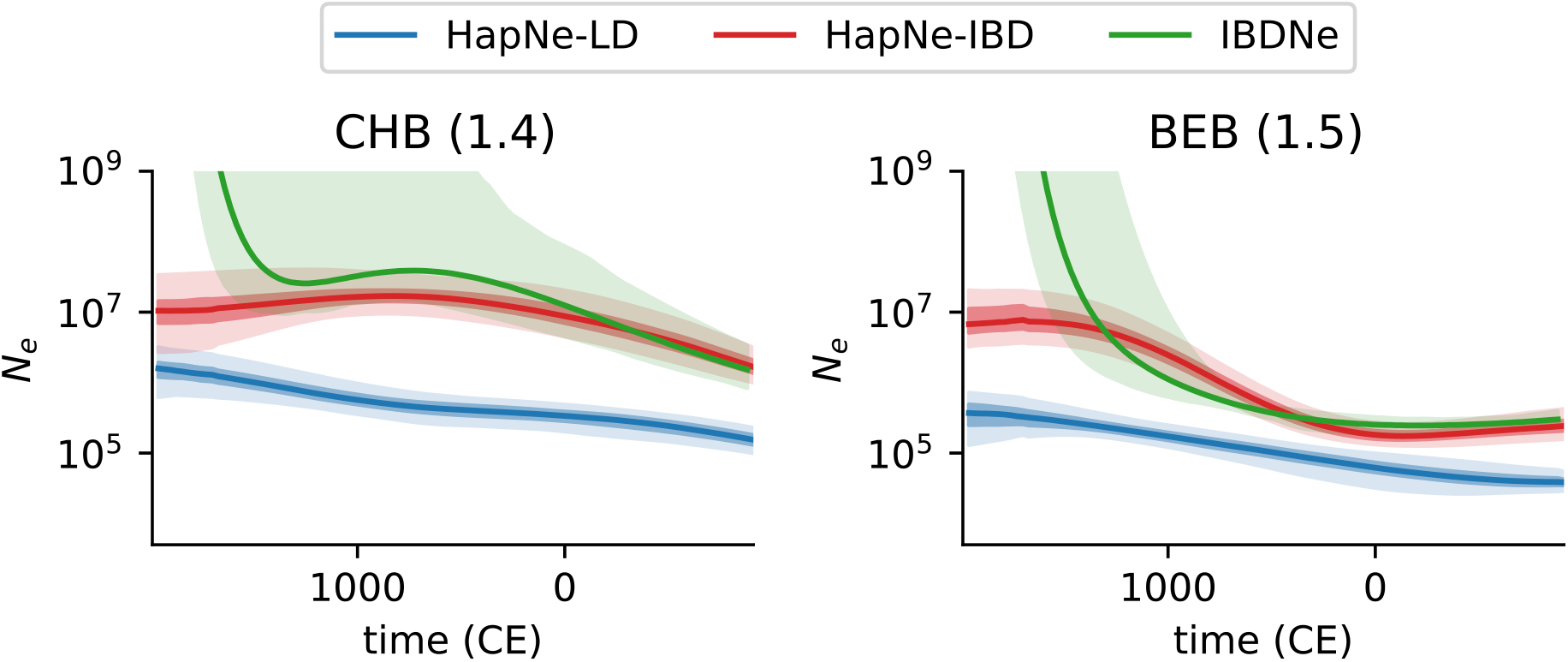
Inferred demographic models for 1,000 Genomes Project populations where significant admixture LD was detected (0.05/26 < *p* < 0.05). Results for populations for which the admixture LD test was significant at 0.05/26 < *p* < 0.05. Numbers in parentheses correspond to – log_10_(*p*). IBD segments for IBDNe and HapNe-IBD were computed using FastSMC.

**Figure S13:**
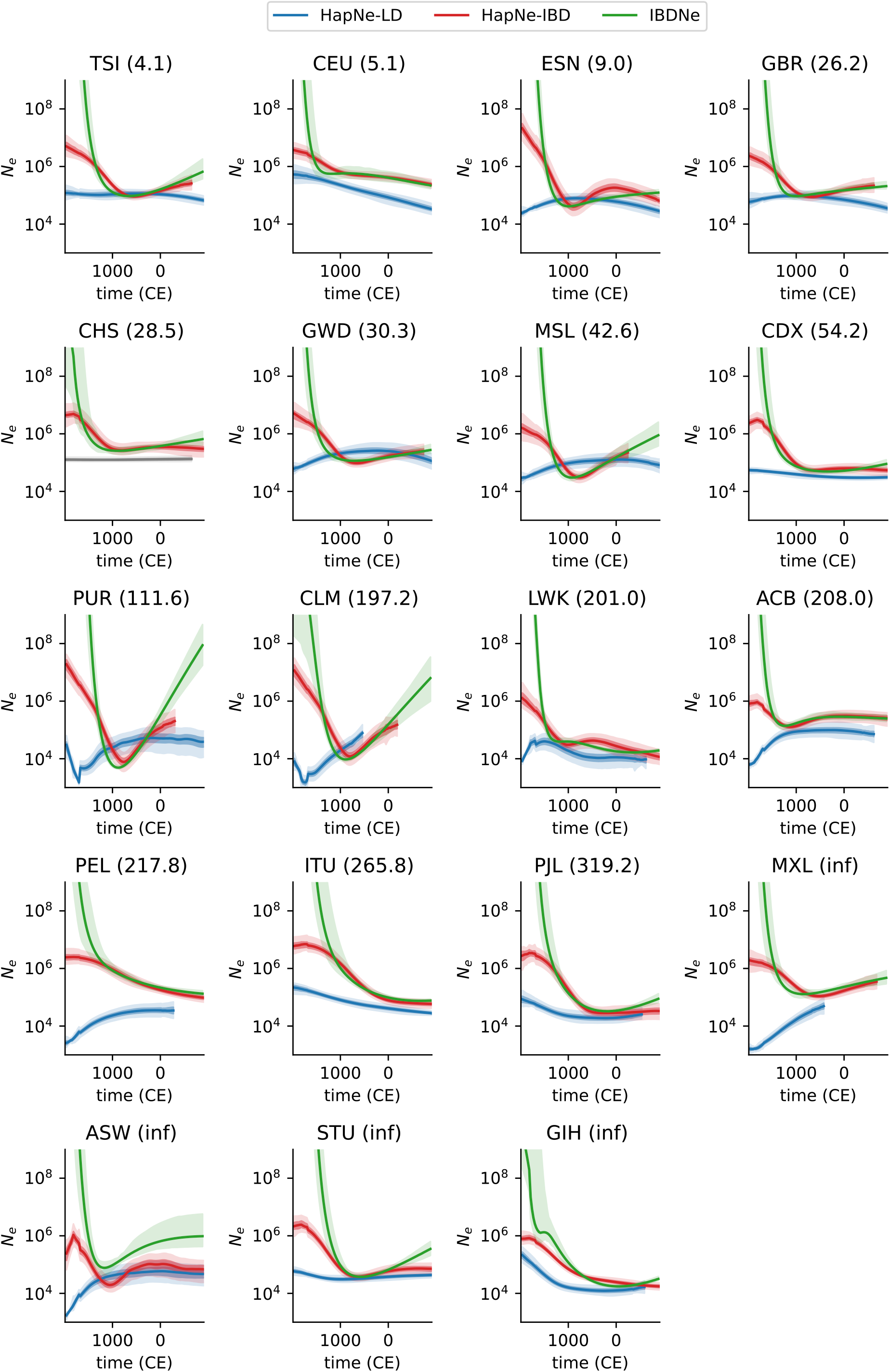
Inferred demographic models for 1,000 Genomes Project populations where significant admixture LD was detected (*p* < 0.05/26). Results for populations for which the admixture LD test was significant at *p* < 0.05/26. Numbers in parentheses correspond to – log_10_(*p*). IBD segments for IBDNe and HapNe-IBD were computed using FastSMC.

#### 1.5 Supplementary Tables

**Table S1:** Further information on populations analyzed in Figure 4. Sample size *s*, average coverage, estimated age of the most recent and distant samples (given in years before 1950), and approximate p-value for the CCLD test for each analyzed ancient population.

| Population | s | Avg. Cov. | Date From (bp) | Date to (bp) | $-\log_{10}$ pval |
| --- | --- | --- | --- | --- | --- |
| Arras in Pocklington | 24 | 2.94 | 2175 | 2202 | 0.54 |
| South England MIA(-LIA) | 49 | 2.88 | 2022 | 2227 | 1.00 |
| Viking Norway | 22 | 1.50 | 950 | 1100 | 1.51 |
| Viking Gotland | 28 | 1.45 | 975 | 975 | 3.52 |
| Caribbean Ceramic | 71 | 2.74 | 510 | 801 | inf |
| Dominican SE coast Ceramic | 18 | 3.08 | 849 | 1150 | inf |

**Table S2:**
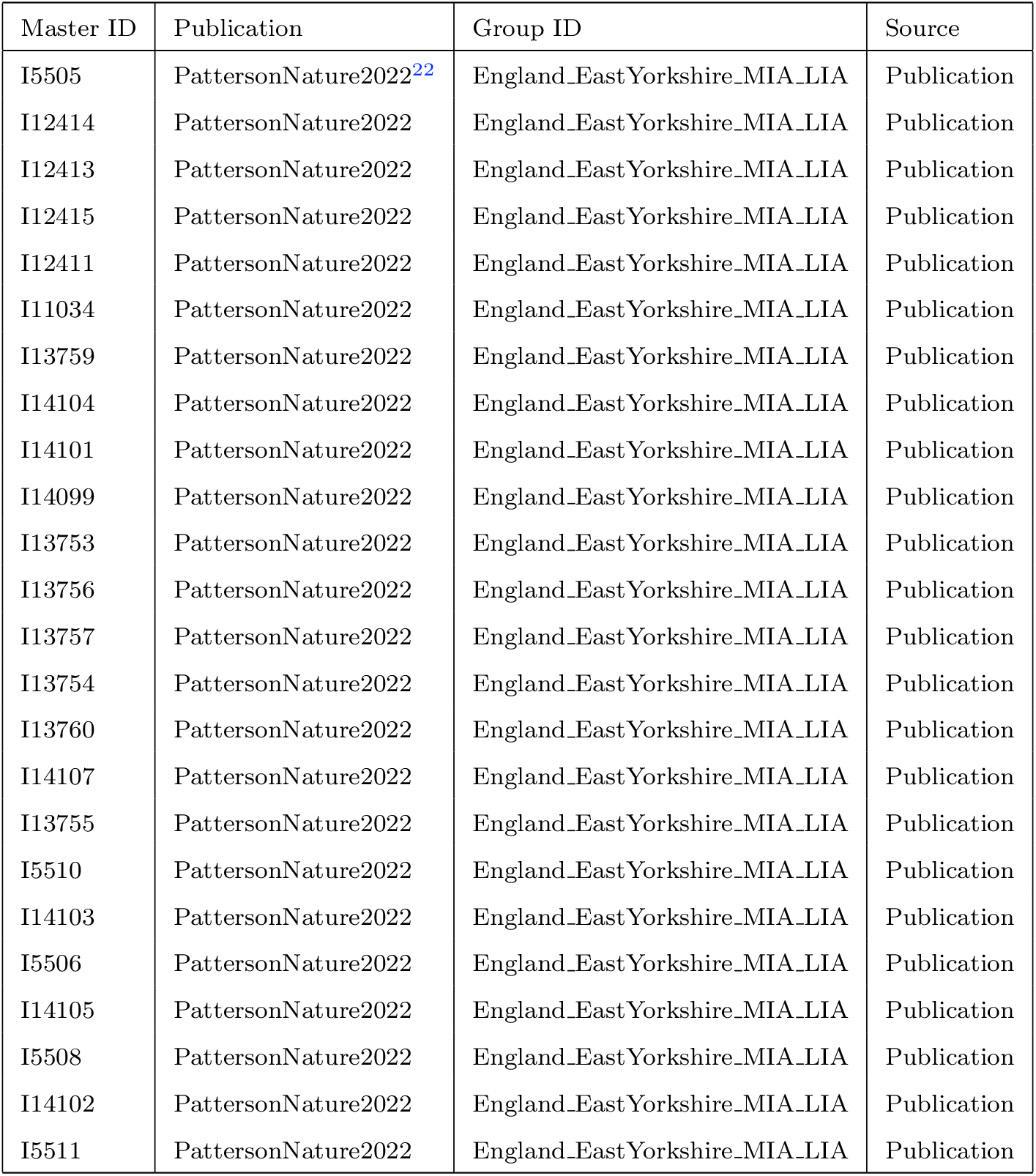
Samples used in the Arras analysis. Genotypes were downloaded from published supplementary materials.

**Table S3:** Samples used in the South England MIA-LIA analysis. Genotypes were downloaded from published supplementary materials.

| Master ID | Publication | Group ID | Source |
| --- | --- | --- | --- |
| I11145 | PattersonNature2022<sup>22</sup> | England_LIA | Publication |
| I19869 | PattersonNature2022 | England_LIA_daughter.I19870 | Publication |
| I16458 | PattersonNature2022 | England_MIA_LIA | Publication |
| I16457 | PattersonNature2022 | England_MIA_LIA | Publication |
| I16450 | PattersonNature2022 | England_MIA_LIA | Publication |
| I17017 | PattersonNature2022 | England_LIA_highEEF | Publication |
| I21308 | PattersonNature2022 | England_MIA_LIA | Publication |
| I11142 | PattersonNature2022 | England_LIA | Publication |
| I27379 | PattersonNature2022 | England_LIA | Publication |
| I21311 | PattersonNature2022 | England_MIA_LIA | Publication |
| I16601 | PattersonNature2022 | England_MIA_LIA | Publication |
| I11992 | PattersonNature2022 | England_MIA_LIA | Publication |
| I21312 | PattersonNature2022 | England_MIA_LIA | Publication |
| I17263 | PattersonNature2022 | England_MIA_LIA | Publication |
| I21310 | PattersonNature2022 | England_MIA_LIA | Publication |
| I11991 | PattersonNature2022 | England_MIA_LIA | Publication |
| I21307 | PattersonNature2022 | England_MIA_LIA | Publication |
| I13726 | PattersonNature2022 | England_MIA_LIA | Publication |
| I11143 | PattersonNature2022 | England_MIA_LIA | Publication |
| I21309 | PattersonNature2022 | England_MIA_LIA | Publication |
| I21313 | PattersonNature2022 | England_MIA_LIA | Publication |
| I20989 | PattersonNature2022 | England_MIA_LIA | Publication |
| I17262 | PattersonNature2022 | England_MIA_LIA | Publication |
| I20987 | PattersonNature2022 | England_MIA_LIA | Publication |
| I20985 | PattersonNature2022 | England_MIA_LIA | Publication |
| I20983 | PattersonNature2022 | England_MIA_LIA | Publication |
| I20986 | PattersonNature2022 | England_MIA_LIA | Publication |
| I20982 | PattersonNature2022 | England_MIA_LIA | Publication |
| I20984 | PattersonNature2022 | England_MIA_LIA | Publication |
| I19657 | PattersonNature2022 | England_MIA_LIA | Publication |
| I19855 | PattersonNature2022 | England_MIA_LIA | Publication |
| I19854 | PattersonNature2022 | England_MIA_LIA | Publication |
| I11993 | PattersonNature2022 | England_MIA_LIA | Publication |
| I11994 | PattersonNature2022 | England_MIA_LIA | Publication |
| I12792 | PattersonNature2022 | England_MIA_LIA_mother.I12793 | Publication |
| I20990 | PattersonNature2022 | England_MIA | Publication |
| I19912 | PattersonNature2022 | England_MIA | Publication |
| I13680 | PattersonNature2022 | England_MIA | Publication |
| I17261 | PattersonNature2022 | England_MIA | Publication |
| I14863 | PattersonNature2022 | England_MIA | Publication |
| I17267 | PattersonNature2022 | England_MIA_LIA | Publication |
| I20988 | PattersonNature2022 | England_MIA_LIA | Publication |
| I17264 | PattersonNature2022 | England_MIA_LIA | Publication |
| I14866 | PattersonNature2022 | England_MIA | Publication |
| I17016 | PattersonNature2022 | England_MIA | Publication |
| I14859 | PattersonNature2022 | England_MIA | Publication |
| I17015 | PattersonNature2022 | England_MIA | Publication |
| I19909 | PattersonNature2022 | England_MIA | Publication |
| I17014 | PattersonNature2022 | England_MIA | Publication |

**Table S4:** Samples used in the Norway Viking analysis. Genotypes were downloaded from V50 of the Allen ancient data resource.^24^

| Master ID | Publication | Group ID | Source |
| --- | --- | --- | --- |
| VK387 | MargaryanWillerslevNature2020<sup>23</sup> | Norway_Viking.SG | V50<sup>24</sup> |
| VK414 | MargaryanWillerslevNature2020 | Norway_Viking.SG | V50 |
| VK530 | MargaryanWillerslevNature2020 | Norway_Viking_o2.SG | V50 |
| VK386 | MargaryanWillerslevNature2020 | Norway_Viking.SG | V50 |
| VK389 | MargaryanWillerslevNature2020 | Norway_Viking.SG | V50 |
| VK393 | MargaryanWillerslevNature2020 | Norway_Viking.SG | V50 |
| VK394 | MargaryanWillerslevNature2020 | Norway_Viking.SG | V50 |
| VK422 | MargaryanWillerslevNature2020 | Norway_Viking.SG | V50 |
| VK515 | MargaryanWillerslevNature2020 | Norway_Viking.SG | V50 |
| VK516 | MargaryanWillerslevNature2020 | Norway_Viking.SG | V50 |
| VK520 | MargaryanWillerslevNature2020 | Norway_Viking.SG | V50 |
| VK524 | MargaryanWillerslevNature2020 | Norway_Viking.SG | V50 |
| VK415 | MargaryanWillerslevNature2020 | Norway_Viking.SG | V50 |
| VK420 | MargaryanWillerslevNature2020 | Norway_Viking.SG | V50 |
| VK448 | MargaryanWillerslevNature2020 | Norway_Viking.SG | V50 |
| VK547 | MargaryanWillerslevNature2020 | Norway_Viking.SG | V50 |
| VK518 | MargaryanWillerslevNature2020 | Norway_Viking_o1.SG | V50 |
| VK392 | MargaryanWillerslevNature2020 | Norway_Viking.SG | V50 |
| VK417 | MargaryanWillerslevNature2020 | Norway_Viking.SG | V50 |
| VK525 | MargaryanWillerslevNature2020 | Norway_Viking.SG | V50 |
| VK526 | MargaryanWillerslevNature2020 | Norway_Viking.SG | V50 |
| VK548 | MargaryanWillerslevNature2020 | Norway_Viking.SG | V50 |

**Table S5:** Samples used in the Gotland Viking analysis. Genotypes were downloaded from V50 of the Allen ancient data resource.^24^

| Master ID | Publication | Group ID | Source |
| --- | --- | --- | --- |
| VK58 | MargaryanWillerslevNature2020<sup>23</sup> | Sweden_Viking.SG | V50<sup>24</sup> |
| VK429 | MargaryanWillerslevNature2020 | Sweden_Viking.SG | V50 |
| VK433 | MargaryanWillerslevNature2020 | Sweden_Viking.SG | V50 |
| VK455 | MargaryanWillerslevNature2020 | Sweden_Viking.SG | V50 |
| VK456 | MargaryanWillerslevNature2020 | Sweden_Viking.SG | V50 |
| VK56 | MargaryanWillerslevNature2020 | Sweden_Viking.SG | V50 |
| VK64 | MargaryanWillerslevNature2020 | Sweden_Viking.SG | V50 |
| VK60 | MargaryanWillerslevNature2020 | Sweden_Viking.SG | V50 |
| VK432 | MargaryanWillerslevNature2020 | Sweden_Viking.SG | V50 |
| VK460 | MargaryanWillerslevNature2020 | Sweden_Viking.SG | V50 |
| VK461 | MargaryanWillerslevNature2020 | Sweden_Viking.SG | V50 |
| VK463 | MargaryanWillerslevNature2020 | Sweden_Viking.SG | V50 |
| VK434 | MargaryanWillerslevNature2020 | Sweden_Viking.SG | V50 |
| VK431 | MargaryanWillerslevNature2020 | Sweden_Viking.SG | V50 |
| VK475 | MargaryanWillerslevNature2020 | Sweden_Viking.SG | V50 |
| VK468 | MargaryanWillerslevNature2020 | Sweden_Viking.SG | V50 |
| VK50 | MargaryanWillerslevNature2020 | Sweden_Viking.SG | V50 |
| VK479 | MargaryanWillerslevNature2020 | Sweden_Viking.SG | V50 |
| VK474 | MargaryanWillerslevNature2020 | Sweden_Viking.SG | V50 |
| VK478 | MargaryanWillerslevNature2020 | Sweden_Viking.SG | V50 |
| VK473 | MargaryanWillerslevNature2020 | Sweden_Viking.SG | V50 |
| VK477 | MargaryanWillerslevNature2020 | Sweden_Viking.SG | V50 |
| VK53 | MargaryanWillerslevNature2020 | Sweden_Viking.SG | V50 |
| VK51 | MargaryanWillerslevNature2020 | Sweden_Viking.SG | V50 |
| VK232 | MargaryanWillerslevNature2020 | Sweden_Viking.SG | V50 |
| VK48 | MargaryanWillerslevNature2020 | Sweden_Viking.SG | V50 |
| VK454 | MargaryanWillerslevNature2020 | Sweden_Viking.SG | V50 |
| VK452 | MargaryanWillerslevNature2020 | Sweden_Viking.SG | V50 |

**Table S6:**
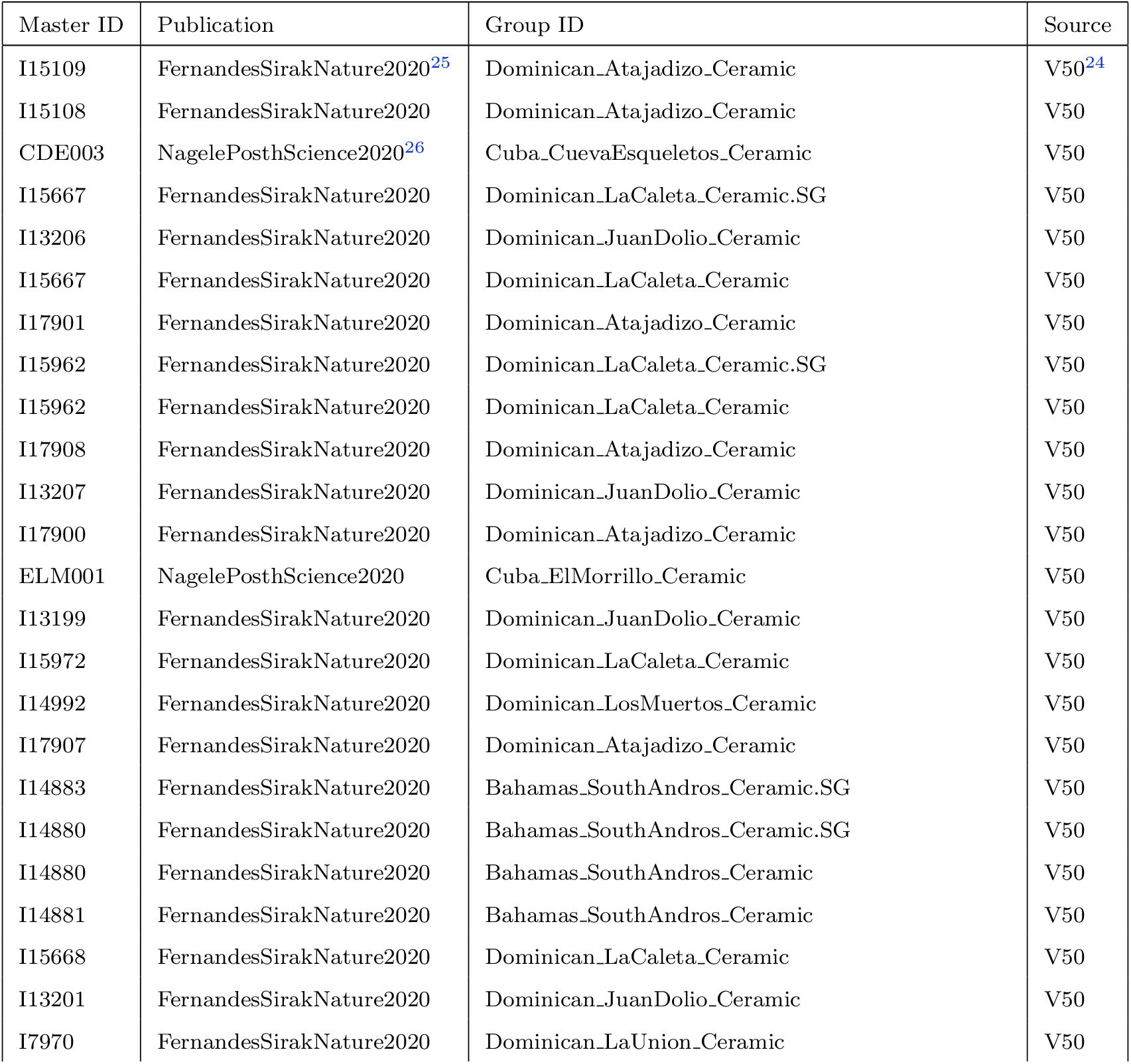
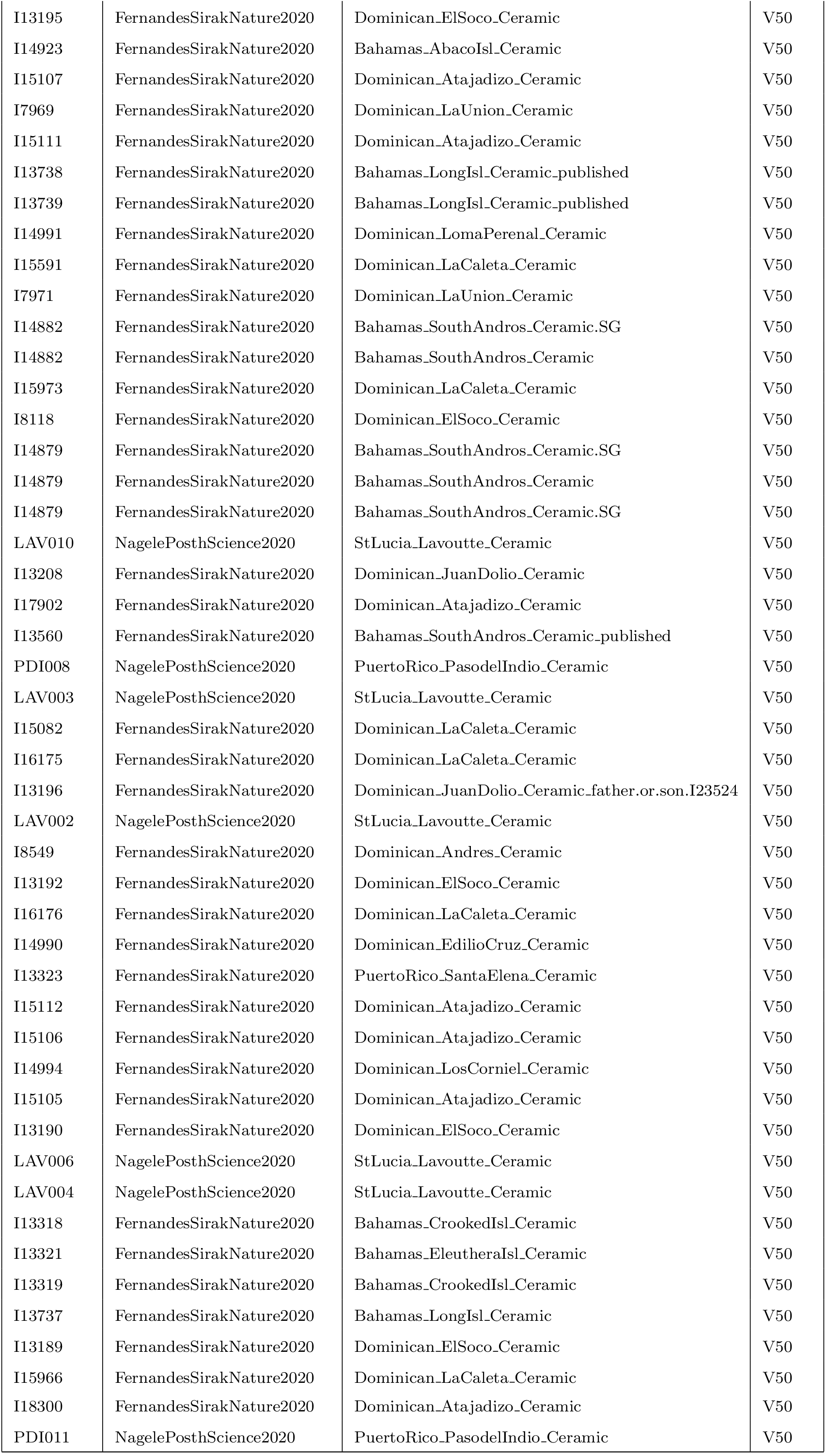
Samples used in the Caribbean Ceramic analysis. Genotypes were downloaded from V50 of the Allen ancient data resource.^24^

**Table S7:**
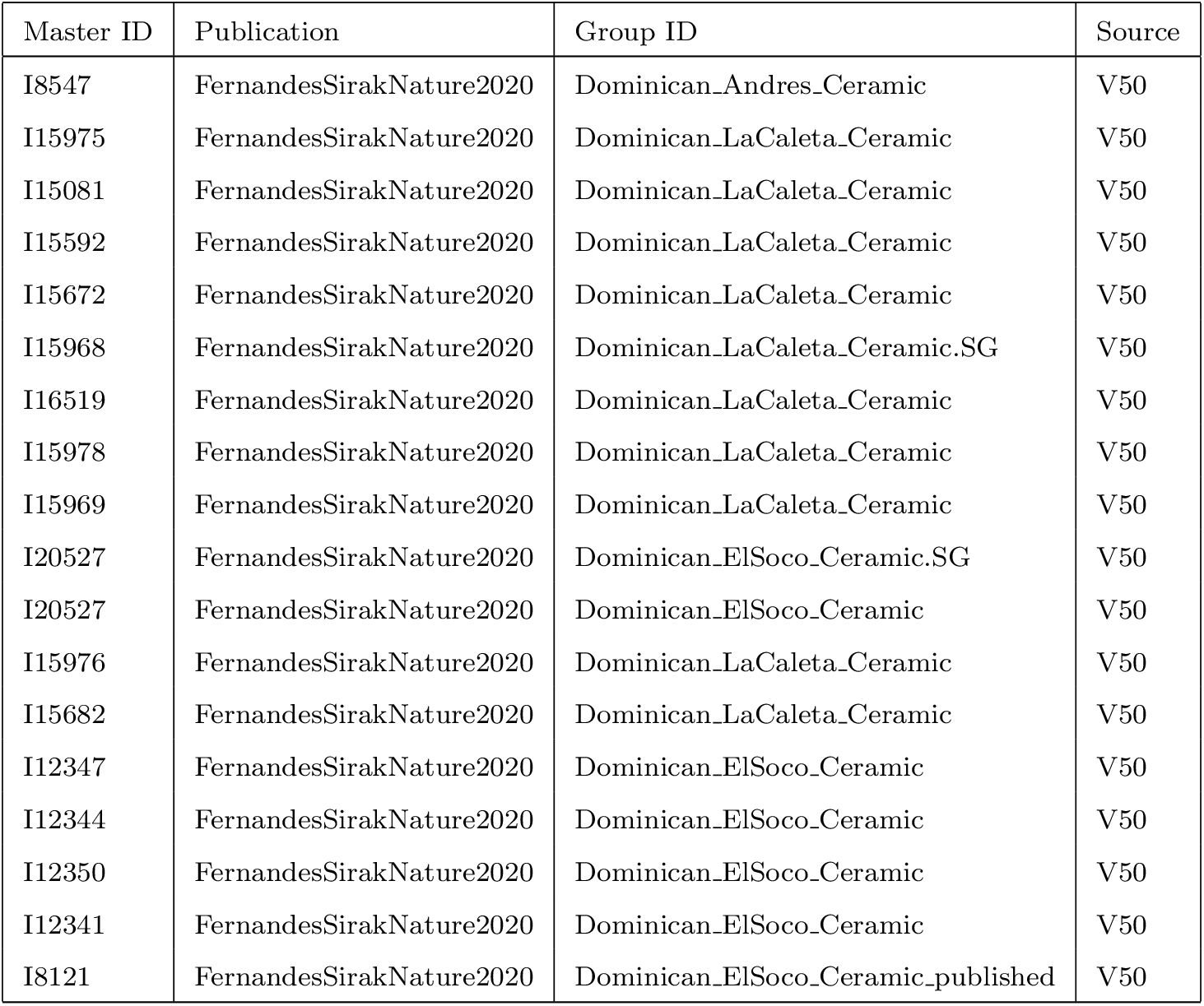
Samples used in the South East Coast Dominican Republic Ceramic analysis. Genotypes were downloaded from V50 of the Allen ancient data resource.^24^

